# Genetic Diversity and Population Structure of the Black-Footed Cat: Insights into *Felis*’s Deadliest Predator

**DOI:** 10.64898/2026.05.29.728895

**Authors:** Victoria B. Grant, Kelsie Hunnicutt, Michelle Schroeder, Martina Küsters, Jonas Oppenheimer, Shreya Banerjee, John J. Baczenas, Dmitri Petrov, Jacqueline M. Bishop, Nadine Lamberski, Beryl Wilson, Alexander Sliwa, Beth Shapiro, Katherine A. Solari, Stepfanie M. Aguillón, Ellie E. Armstrong, Molly Schumer

## Abstract

**Background:** Black-footed cats (*Felis nigripes*) are one of Africa’s least studied felines. The population dynamics and demographic history of this solitary species have not been well-described. Reports of ongoing decline of present-day populations resulted in the IUCN Red List categorizing the species as vulnerable to extinction. As populations decline and become isolated from each other, they become susceptible to strong genetic drift and inbreeding, which can lead to the accumulation of deleterious alleles and increased sextinction risk. However, the IUCN cited data deficiencies across the species range as a limitation in this categorization for black-footed cats. In cases where ecological surveys are lacking, range-wide population genomic surveys can improve our understanding of population dynamics.

**Results:** In the first genomic study of free-roaming individuals, we sequenced whole genomes of black-footed cats (N=44) from across their distribution. To do so, we incorporated whole genome sequences generated from both modern biological samples and century-old museum specimens. We assembled a highly contiguous reference genome using a combination of PacBio HiFi data and publicly available Hi-C data and investigated the demographic history, population structure, and genetic diversity of wild black-footed cats. We found evidence of historical effective population sizes of ∼11,500 individuals, which is lower than estimates reported in other felid species. Consistent with modest historical population sizes, we found that present-day genome-wide diversity was low (*π* ≈ 0.0004). However, despite low genetic diversity, we find that black-footed cat genomes do not harbor long runs of homozygosity. Simulation results indicate that low present-day genetic diversity may simply result from modest historical population size. However, other analyses point to evidence of a population contraction in the last 50 generations, which could contribute to future genomic erosion. We also compared genomic variation in populations across the range to evaluate patterns of population structure, finding evidence of higher genetic similarity between individuals in closer geographic proximity.

**Conclusion:** Overall, these results provide range-wide information about the demographic history and present-day genetic diversity of an understudied species. Together with analyses of population structure, we speculate that there may be greater connectivity between populations of black-footed cats than previously assumed. Our study underscores the utility of genomic data in providing insights into population dynamics for better conservation management.

## Introduction

The threat to biodiversity posed by anthropogenic change has gained global concern as flora and fauna across the world continue to disappear at alarming rates. Vertebrates alone show an average decline in abundance of ∼25%, with scientists increasingly describing this decline as the sixth mass extinction event^1^. As populations decline, both the total number of individuals and the genetic diversity present across all populations decreases^2^. Small, isolated populations are susceptible to the fixation of deleterious alleles due to strong genetic drift and inbreeding, which can result in a negative feedback loop known as the “extinction vortex”^3,4^. This occurs because strong drift increases the likelihood that beneficial alleles will be lost from the population and that deleterious alleles will be fixed, contributing to reduced average fitness^5,6^. Moreover, decreases in standing genetic variation may limit species’ ability to adapt to rapidly changing environments which is a growing concern in the context of human-induced habitat disruption and climate change. Genomic data can help us uncover crucial evolutionary processes affecting contemporary population dynamics to better understand species decline.

Rapid advances in whole genome sequencing and plummeting costs have made genomic approaches more accessible to researchers focusing on understudied organisms in a conservation context^7–9^. As a result, this data can be harnessed to study non-model organisms’ present-day genetic diversity and evolutionary past. Traditional conservation genetic techniques utilize putatively “neutral” genetic markers to investigate population dynamics. The availability of genome-scale data dramatically increases the number of markers that can be used to address these questions^8^, enabling researchers to infer additional information like migration rates or evidence of loss of adaptive variation^8,10–12^.

Despite growing recognition of the influence of genetic diversity and population history in species resilience^13,14^, genomic data has not been broadly incorporated into conservation assessments. The IUCN Red List acts as the global database for species’ extinction risk but still relies on approximate methods to estimate the census population (N), or the number of reproductive individuals within a species, to inform policy decisions. However, while N summarizes current population size and may be useful in assessing immediate extinction risk, past demographic events like population bottlenecks can have disproportionate impacts on contemporary genetic diversity and the number of ancestral lineages represented in present day individuals. Thus, the effective population size (N_e_) is often a useful measurement summarizing the number of individuals consistent with the population genetic properties of the present-day population. N_e_ captures the effects of past demographic events on genetic variation and places patterns of present-day genetic diversity in the context of historical population dynamics^15–17^. Thus, inferences of demographic history, effective population size (N_e_), and population structure inferred from genomic data can be leveraged to improve population risk assessments and management strategies for species of conservation concern^10^.

Black-footed cats (*Felis nigripes*), one of only three endemic African *Felis* species, are one of the continent’s rarest cats with the most restricted species distribution of African felids (Figure 1A & B)^18,19^. Black-footed cats belong to the domestic cat lineage (*Felis catus,* Figure 1C) and are carnivores who operate as keystone predators at their trophic level. This enigmatic species has gained recognition for their incredibly high hunting success rate (∼60%), and have been described as the deadliest solitary hunter in popular media^20^.

**Figure 1.**
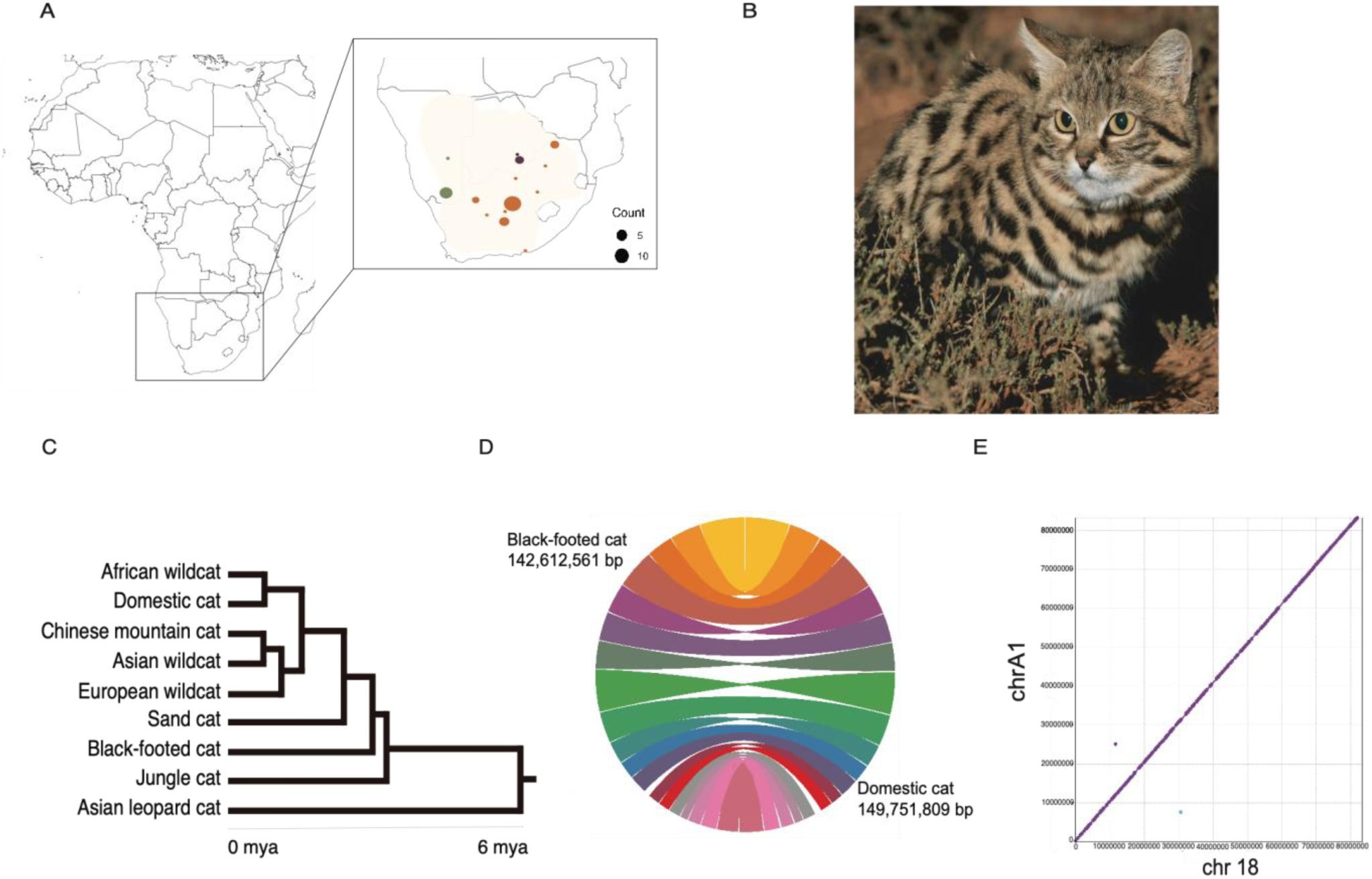
Sample collection map, comparison between black-footed cat and domestic cat assemblies and *Felis* phylogeny. (**A)** Distribution of black-footed cat samples collected for this study throughout the species distribution range (shaded area). Points indicate approximate sampling locations and do not reflect exact capture sites, some of which are unknown. Colors correspond to country of origin (Botswana – eggplant, Namibia – olive, and South Africa – orange). Point size corresponds to the number of samples that originated from that site in our dataset. **(B)** An adult, male black-footed cat from the De Aar site, South Africa population. Photo credit to A. Sliwa. **(C)** *Felis* phylogeny adapted from Li et al 2024. **(D)** Circos plot comparing genome assemblies of the black-footed (this study) and domestic (Buckley et al 2020) cat. The 19 chromosomes in the black-footed cat genome assembly are depicted on the left side and those of the domestic cat (FelCat9) are depicted on the right side. Each color in circos plot corresponds to a different chromosome. Scaffold N50 for each species’ assembly is noted under their respective names. **(E)** Mummer alignment between chromosome A1 assembled in this study and its homologous chromosome from another high-quality black-footed cat assembly (Yuan et al 2024). Purple indicates co-linear alignments and blue indicates flipped alignments. Note the high concordance between these two long-read assemblies.

Based on monitoring efforts, black-footed cats appear to have undergone a steady population decline and are listed as vulnerable to extinction on the IUCN Red List^19^. Evidence for their vulnerable status stems primarily from the long-term study population in the Benfontein Nature Reserve in Kimberley, South Africa. Black-footed cats occur in low densities with occurrences of ∼0.03 cats per km^2^ at peak estimated densities, making census population size difficult to establish. The study area in the Benfontein Nature Reserve is a protected space known to have higher densities of black-footed cats, so the decline in this area from 1998-2015 and increased natural mortality could indicate population decline in the species overall^19,21^. Population declines have been attributed to anthropogenic pressures including habitat loss, prey availability, and predation^22,23^. Changes in habitat driven by human development, and further disruption anticipated from climate change, have spurred concerns about the impacts of population fragmentation and potential inbreeding.

The most recent IUCN Red List assessment notes data deficiencies on the black-footed cat, including a lack of systematic field surveys across the species range and lack of research on geographically distinct subpopulations, that may limit their assessment of extinction vulnerability. Moreover, efforts to maintain black-footed cats in captivity, which has been beneficial in conservation efforts in other species^24–26^, have been largely unsuccessful. In captivity, their typical ∼5-6 year lifespan is reduced to ∼2-5 years, and they experience high mortality, often caused by an excess buildup of amyloid protein, known as amyloidosis^27^. The difficulty of maintaining black-footed cat populations in captivity makes conserving genetic diversity in the wild even more crucial. The only previous genomic study on this species used zoo and museum specimens and found evidence of low genetic diversity in black-footed cats^28^. However, this study only had access to museum samples from captive populations of unknown geographic origin. Genomic data from natural black-footed cat populations will allow for analyses of population structure, connectivity, and genetic diversity in this understudied group.

Despite broad concerns about extinction risk, black-footed cats’ small size and nocturnal behavior contribute to them being one of Africa’s least-studied felids. Little is known about their evolutionary history, genetic variation, or whether genetically distinct subpopulations exist. Here, we present results on whole genome sequencing data from 44 individuals sampled across their geographic range, representing the first genomic study of free-roaming, wild black-footed cats. Our dataset incorporates whole genome sequences generated from biological samples from three geographic regions as well as preserved specimens that stretch back to 1818. We analyze this data with the goal of addressing gaps in our knowledge about black-footed cats’ demographic history, inbreeding risk, and present-day population structure. To this end, we produced a highly contiguous reference genome using PacBio HiFi sequencing and publicly available Hi-C data^29^. Together, we find evidence of low genetic diversity across the black-footed cat range, likely attributable in part to modest historical population sizes. We also report surprisingly weak population substructure across geographically distinct populations, hinting at higher than expected levels of connectivity between within-country populations or more recent dispersal events. Our study provides valuable insights into black-footed cat evolutionary dynamics that can improve our understanding of extinction risk in this endemic species.

## Methods

### Sample collection

Black footed cat blood and tissue samples (N=44) were collected from a variety of sources with the aim of covering locations across the species range (Figure 1A). Samples were provided by the Black-footed Cat Working Group (BFCWG), primarily from their main study site in Kimberley, South Africa at the Benfontein Nature Reserve (hereafter Benfontein) established in 1992. Since 2004, the group has also periodically conducted opportunistic sampling throughout South Africa. Benfontein is a private nature reserve stocked with wild game species. It harbors diverse vegetation in an open area and experiences heavy rainy seasons (average 450 mm rainfall annually). Blood samples were taken from black-footed cats caught during radio collaring expeditions and frozen. Tissue samples were collected opportunistically from individuals killed in vehicular collisions throughout the Northern Cape Province of South Africa. Samples from these expeditions were frozen and stored at the San Diego Wildlife Alliance, USA. For several samples, cell pellets were developed at the San Diego Wildlife Alliance and stored at -80°C before being provided for this study.

Samples from Namibia were provided by a project of the BFCWG, the Black-footed Cat Research Project Namibia (BFCRPN). The BFCRPN was established in 2012 to initially collect distribution records and identify areas for ecological studies in Namibia. The Grünau black-footed cat study site was established in 2020. In contrast to Benfontein, Grünau has a much more arid climate in the Nama Karoo with little rain and common droughts (80-100 mm rainfall annually). The Grünau site is also located on farmland, with less diverse vegetation, inhabited by moderate densities of sheep. Hair samples were collected during radio collar expeditions in Grünau (CITES #0067783).

Samples (N=27) collected from South Africa and Namibia by the BFCWG and BFCRP were collected over the past 20 years and are considered modern samples for the purpose of extraction, library preparation, and sequencing (referred to as non-preserved below). In addition to the modern samples, samples taken from mounted specimens (N=17) were collected from natural history museums within the United States and South Africa, and additionally from private collections in Namibia. Given their age, these samples required a distinct DNA extraction and library preparation protocol to account for DNA degradation.

### DNA extraction and library preparation for non-preserved samples

For all non-preserved samples, we used the QIAGEN DNeasy Blood and Tissue Extraction Kit on the blood, tissue, and cell samples following the manufacturer’s specified protocol for the appropriate sample type. Based on the resulting DNA concentration measured on a Qubit 4 fluorometer, we varied our library preparation approach.

For extractions that resulted in high DNA yields from samples (≥15 ng/µL), we used a protocol adapted from Quail^30^. At least 40 µL of DNA was sheared to 350-450 basepairs length using a Qsonic sonicator. After DNA was sheared to the desired length, 12.5 µL of an end-repair mastermix (DNA ligase buffer, 10mM dNTP, T4 DNA polymerase, Klenow DNA polymerase, T4 polynucleotide kinase) was added to 37.5 µL sonicated DNA then incubated at room temperature for 30 minutes. After incubation, the DNA end-repair mix was purified with the QIAGEN QIAquick PCR purification kit before 11.5 µL of the A-tailing mix (containing Klenow buffer, dATP, and Klenow exonuclease) was added then incubated again for 30 minutes in a thermomixer at 37°C. We repeated purification then added 7.5 µL of adapter ligation mix (ligase buffer, paired-end adapter mix, DNA ligase) before the final incubation at room temperature for 15 minutes.

The mix was purified again with the QIAGEN QIAquick PCR Purification Kit then quantified using a Qubit 4 fluorometer. We amplified the sample with 14 µL Phusion PCR mix (HF buffer [NEB, B0518S], custom index primer, dNTPs [N0447S], Phusion High-Fidelity DNA Polymerase [NEB, M0530]) for 12 cycles. After the PCR run, we purified the library using Agencourt AMPure XP beads following the manufacturer’s instructions. Final library size distribution and concentration was characterized on the Agilent TapeStation and Qubit.

Some of our samples were DNA aliquots previously extracted using an unknown protocol. After quantifying these samples with a Qubit, we found these extractions tended to have low DNA concentrations (9-15 ng/μL) compared to the other modern samples we extracted using the QIAGEN DNeasy Kit. (11-260 ng/μL). For all extractions with concentrations below 15 ng/μL, we used an in-house protocol for low-coverage whole genome sequencing with a tagmentation enzyme. We first diluted the DNA with water to 1.5ng/μL and used 1.5 μL of sample. We added 1.5 μL of Tagment DNA buffer and Tagment DNA Enzyme mixture from Illumina Tagment DNA kits (20034197) to each sample before samples were centrifuged for 5 minutes at 55°C. We then added Kapa Master Mix, the i7, and i5 index for each sample before we placed them in the thermocycler for PCR amplification (72℃ for 3 minutes, 98℃ for 2 minutes and 45 seconds, 8x of 98℃ for 15s, 62℃ for 30s, 72℃ for 90s).

Following amplification, we briefly spun down the tubes and added 18 µL of silica beads to each library. We let libraries sit for 5 minutes at room temperature before we separated them on the magnet for 1 minute. We discarded the supernatant then washed the pellet with 50 µL of 70% ethanol while on the magnet. After we repeated the ethanol wash, we air dried tubes for 5 minutes and added 15 µL of TrisHCl to resuspend the beads. Tubes sat at room temperature for 2 minutes, were placed on the magnet for 1 minute, and we then moved the purified library into Eppendorf LoBind tubes.

### DNA extraction and library preparation for preserved samples

Mounted specimens were included in this study to access more individuals across the species distribution. These mounted specimens came from both private and museum collections and were preserved through a variety of methods including sun-tanning and salt baths. For mounted specimen hair and tissue samples, 1 cm^2^ subsamples were collected from previously preserved specimens. We prepared the subsamples for extraction by performing a series of washes on the subsampled tissue. First, tissue samples were washed in two UltraPure H20 water baths while hair samples were washed in one 0.5% bleach solution and three water baths. After being washed, samples were chopped with a sterile scalpel blade before digestion. We digested samples in 11 µL of Gilbert et al 2007^31^ lysis buffer (10% SDS, Ultra Pure H20, 1M Tris, EDTA, 5M NaCl, 1M CaCl2, 1M DTT, Proteinase K), without UV radiation, overnight (15-24 hours) on a shaking incubator (1250 rpm) at 55°C then proceeded with the Rohland et al extraction^31^, including a negative control.

The digested tissue was removed from the shaker and placed into an ice bath to cool for 5 minutes then pelleted by microcentrifugation at 15,000 rpm for 2 minutes. Prior to starting DNA isolation, we resuspended the silica bead stock solution (G-Biosciences, 786-915) by vortexing. Silica beads were placed on a magnet where supernatant was pipetted off for two 80% ethanol washes before final resuspension in the EBT elution buffer. We transferred 1.56 mL binding buffer D and 10 µL of washed silica beads to Eppendorf LoBind tubes. Avoiding beads, we added 150 µL of the lysate to the binding buffer/bead solution, vortexed, and rotated at room temperature for 15 minutes. After rotation, we spun the solution into a pellet, placed on a magnet for 1 minute, then discarded the supernatant. We then added 250 µL buffer PE, vortexed the sample, and spun back into a pellet. The pellet was placed back on a magnet for 1 minute then the supernatant was pipetted off and discarded. We repeated the buffer PE wash for a total of three washes before we dried the beads with open lids on a magnet for 20 minutes at room temperature. We then added 14 µL of buffer EBT and vortexed before we spun the sample to collect the beads. The samples were incubated at room temperature for 5 minutes. After incubation, we transferred the supernatant to a fresh Eppendorf LoBind tube and repeated buffer EBT elution for a final DNA volume of 28 µL.

To prepare libraries for sequencing from these potentially degraded samples, we used a single-stranded library preparation method designed for ancient DNA extractions^32,33^. We combined 10 µL of DNA extract with 1µL of 76 ng/µL SSB (NEB, M2401S) in PCR tubes. We denatured input DNA in the thermocycler at 95°C for 3 minutes with a lid temperature of 105°C. Then, we cold shocked the denatured DNA on an ice block for 2 minutes before we pipetted 1 µL of 2 µM annealed P5 adapter/splint and 1 µL of 0.4 µM P7 adapter/splint. We added 12 µL of mastermix to each tube before the sample was vortexed and incubated in a thermocycler at 37°C for 1 hour. After incubation, we added 35 µL of EBT buffer and 60 µL 18% PEG SPRI to each reaction followed by two 180 µL 80% ethanol washes. We then eluted reactions in 20 µL EBT buffer after two washes.

To determine the number of PCR cycles necessary for library amplification, we quantified the libraries using qPCR mixing 1X Maxima SYBR Green PCR Master Mix with 200 nM IS7 and, 200 nM IS8 primers into 2µL dilution of 1:100 library. Based on qPCR Ct values, we performed 9-15 PCR cycles using 1X Amplitaq Gold 360 Master Mix and unique dual index, 2uM i7 index primer and 2uM i5 index, primer combinations at 95°C for 10 minutes, 95°C at 30 seconds, 60°C for 30 seconds, 70°C for 1 minute, then 72°C for 7 minutes, for the samples based on recommendations from the Spotlight v1.2.2 2020 library preparation protocol^33,34^. We cleaned the amplified library using a SPRI bead-based purification protocol.

### Sequencing

Libraries were pooled together in batches of approximately 7 libraries per lane and sequenced on an Illumina NovaSeq 6000. The modern samples, containing libraries from high and low DNA concentration extractions, were sequenced to a target read depth of ∼450 million 150 bp paired end reads per sample. Libraries from preserved specimens required higher read depth to achieve similar coverage due to quality differences in libraries. Prior to deep sequencing, we evaluated endogenous content, mapping rate, and library complexity through a small sequencing run to a depth of ∼1 million reads. We sequenced preserved libraries that had over 70% endogenous content, >80% mapping rate, and at least 80% library complexity to a target depth of ∼900 million reads.

We additionally chose to sequence a subset of preserved samples (N=3) from geographically distinct locations, which happened to be museum specimens, to be sequenced to a final depth of ∼10X on one lane of the NovaSeq 6000 platform (150bp, paired end). Coverage per sample can be found in Table S1.

### Reference genome preparation and assembly

A cell sample from one male individual (id #15015) from the San Diego Wildlife Alliance containing ∼2×10e6 cells was selected to generate PacBio HiFi data for a reference genome. This sample was collected from a free-roaming South African individual and later stored at the San Diego Wildlife Alliance. We conducted a high molecular weight extraction through the Stanford Genomic Sequencing Services Core. The extracted DNA was processed at UC Berkeley QB3 for PacBio single-molecule real-time (SMRT) library preparation and sequenced on 3 SMRTcells of the Sequel II platform.

We used hifiasm v0.18.2^35–37^ with default settings to assemble a draft genome for this sample. In order to improve the contiguity of the genome assembly, we used publicly available Hi-C data from DNAZoo (SRR13167949) to scaffold the primary assembly. We followed the recommended instructions from YaHS^38^ in order to prepare the genome for scaffolding. Briefly, we used BWA v0.7.17^39^ index with flags ‘-a bwtsw’ to index the primary assembly. Forward and reverse Hi-C reads were mapped independently using BWA-mem^40^, and 5’ ends subsequently filtered using scripts from YaHS v1.1^38^. Reads were then paired and combined into a single BAM file and reads with MAPQ less than 10 were filtered out using SAMtools v1.16.1^41,42^. Reads were then sorted and BAM files indexed using SAMtools^41^. Finally, read groups were added and duplicates removed using Picard Tools^43^ AddOrReplaceReadGroups and MarkDuplicates. Further details can be found on the Arima Genomics mapping pipeline GitHub (https://github.com/ArimaGenomics/mapping_pipeline). Using YaHS^38^ with default parameters, we then generated a scaffolded assembly in addition to contact maps for visualization in Juicebox^29^. Juicebox input files were generated using the recommended instructions on the YaHS^38^ GitHub (https://github.com/c-zhou/yahs) using a combination of Juicer Tools^29,44^ and Juicer^29^. We then imported the relevant files into Juicebox and made several visual joins. Finally, we decontaminated the assembly using the NCBI FCS-GX pipeline^45^.

In order to investigate the quality and contiguity of our assembly, we summarized assembly statistics (e.g. N50, L90, number of scaffolds/contigs; Table S2) and used BUSCO^46,47^ analysis to evaluate gene completeness in comparison with other genomes from the *Felis* genus (Table S3). Finally, we produced a whole-genome alignment of our assembly with the domestic cat (Felcat9; GCA_000181335.4) in order to identify autosomal scaffolds and look broadly at synteny (Figure 1D)^48^. We used scripts from Assemblathon2^49^ to evaluate basic assembly statistics and subsequently ran BUSCO v5.4.4 using both the mammalia_odb10 and carnivora_odb10 dataset (Table S4)^46,48^. We selected five resequenced black-footed cat individuals, with known sex information, to map to the draft assembly in order to identify putative sex chromosomes based on coverage.

To compare our genome assembly with the Yuan et al 2024^28^ genome assembly, we aligned the chromosomes from each assembly to each other to assess contigs using MUMmer^50^. MUMmer^50^ aligns genomes to each other to detect similarity or differences in sequences^50^. In MUMmer^50^, we indexed genomes with *numcer* before using *delta-filter* to align. Alignments were visualized in MUMmer^50^ with the *mummerplot* command. Alignments showed high synteny across all chromosomes. A representative alignment is shown in Figure 1E.

### Assembling the mitochondrial genome and removing potential nuclear mitochondrial DNA segments

In addition to the nuclear genome, we wanted to include a mitochondrial reference haplotype in our assembly. This is particularly important for felid lineages because of known non-functional mitochondrial DNA segments that translocated into the nuclear genome sometime in the evolutionary history of the lineage^51^. These sequences can be associated with high error rates in nuclear variant calling due to mismapping of mitochondria reads to nuclear sequences or vice versa^52^. We separately assembled the mitochondrial genome using MitoHiFi v3.2^53^ using the complete black-footed cat mitochondrial genome^54^ from NCBI (NC_028309.1) as a guide. We next used BLAST^55^ to identify partial mitochondrial sequences present as short scaffolds (<1 Mb) in the reference assembly and removed them. Finally, we generated a new version of the genome that included the assembled mitochondria.

### Bioinformatic processing of modern and historical sequencing data

For modern samples, we performed an initial quality assessment of FASTQ files using FASTQC^56^. For libraries that passed quality assessments, we trimmed adapters from sequences using Cutadapt v. 1.18_py36^57^. We mapped the trimmed reads to our reference assembly using BWA-mem v. 0.7.17^39^ and used SAMtools^41^ to sort the resulting SAM files and generate BAM files.

We summarized the mapping results with SAMtools^41^ *flagstat* and *depth* commands to determine genome depth and average coverage for each sample file. We followed by marking and removing PCR duplicates within BAM files using Picard^43^ *MarkDuplicates.jar* with the *REMOVE_DUPLICATES* flag activated^43^. We found a high percentage of reads mapped across the genome (percent basepairs covered >97%), yielding ∼7-32X coverage per individual (Table S1).

Historic samples required additional filtering before alignment due to smaller fragment size, lower library quality, and non-endogenous sequences. We used Paleomix^58^ 1.3.8 to trim adapter sequences from raw reads with AdapterRemoval v. 2.2.0^58,59^. Within Paleomix^58^, we then used BWA^39^ v. 0.7.15 to map trimmed sequences to the black-footed cat reference genome and generated BAM files including aligned reads with a mapping quality score above 20^39^. Mapped sequences were indexed with SAMtools v. 1.3.1^41^ before being processed with Picard Tools^43^ v. 1.137 *ValidateSamFileto* which generated final BAM files. PCR duplicates were filtered using the *MarkDuplicates* pipeline from Picard Tools^41,43^. Paleomix^58^ reports quality metrics for each BAM file including map damage, genome coverage, and depth which were later used to assess the quality of the alignment. Additionally, we checked the BAM files with SAMtools^41^ *flagstat* and *depth* commands to determine genome depth and average coverage for each sample. BAM files generated using Paleomix^58^ varied in coverage from 1-10X (Table S1).

With both modern and historic BAM files, we ran BCFtools^42^ v. 1.16 *mpileup* command with the ‘--fasta-ref’ and ‘--skip-indels’ flag to generate genotype likelihoods for both variant and invariant sites without indels which were outputted in VCF format using ‘--output-type’. Genotype likelihoods were piped into BCFtools^42^ *call* with ‘--multiallelic-caller’ and again outputted into VCF format using ‘--output-type’. We then indexed the VCF using SAMtools *tabix* before filtering to keep autosomes using the BCFtools^42^ *view* ‘--regions’ flag with the same output flags previously used. From our reference assembly, we identified low mappability and repeat regions and output these regions into a BED file. Using this file, we removed these regions with bedtools v2.17.0^60^ *intersect* accompanied with the ‘-a’ and ‘-b’ flags to identify input as VCF and BED format, respectively, followed by the ‘-v’ flag to report ‘a’ input that does not overlap with ‘b’ input.

Historical samples are susceptible to DNA damage which is often detectable as a bias in the types of variants detected, specifically an excess of transitions^61^. Transition bias is estimated by taking the ratio of sites that are transitions (ti) or transversions (tv) compared to the reference genome. Departures from this expected ratio (ti/tv of ∼2 in mammals) can indicate an excess of errors, and skew interpretation of results^62,63^. To assess potential transition bias, we calculated the transition ratio in our filtered VCF, which included only biallelic SNPs on autosomes. We first masked repetitive regions using with VCFtools^64^ ‘--filter-summary’. We did not find evidence of excess transversions in our transition to transversion ratio (ti/tv=1.4)^65,66^.

### Downsampling BAM files for similar coverage

Differences in coverage have been shown to influence the results of population genetic analyses^67^. To prevent this, for a subset of analyses we generated BAM files that were downsampled to have similar coverage across samples using SAMtools^41^. We indexed BAM files with SAMtools^41^ *index* before we counted mapped reads in each BAM file. We then reduced the number of reads for each BAM file with *view* ‘--bam’ and ‘--S’ to subset the data to 1X genome-wide coverage for each sample (Table S5). We confirmed coverage was reduced to the desired level with SAMtools^41^ *depth*. We generated additional downsampled BAM files for two high coverage, preserved individuals from geographically distinct locations (sample ids 118484 and 6494) to normalize coverage among these samples for a subset of analyses at matched coverage (∼7X coverage).

### Population structure across the species distribution

We performed principal component analysis^68,69^ (PCA) to evaluate genetic differentiation between individuals and infer the presence of population structure. We visualized our results using ggplot2^70^ and conducted a Pearson’s correlation test, using the *cor.test()* command, to test for any correlations between resulting clusters and coverage ^71^. Based on initial analyses that highlighted a strong impact of coverage on clustering, we used samples downsampled to 1X coverage (see Methods and Supporting Information S1). We used ANGSD v. 0.931^72,73^ ‘-GL 1’ flag to generate genotype likelihoods at each SNP for individual samples in geno file format. We filtered the geno file with ‘*-*minMapQ’ and ‘-minQ’ flags for a respective minimum mapping quality and minimum base quality of 20 then implemented the ‘-SNP_pval’ flag for a p-value threshold for calling SNPs of 0.001 and removed transitions with the ‘-doGeno 1’ flag. Non-biallelic sites were excluded when generating the geno file and we only analyzed uniquely mapping and properly paired reads with ‘-uniqueOnly 1’ and ‘-only_proper_pairs 1’ flags. Additionally, we calculated base alignment quality with ‘-BAQ 1’, major and minor allele frequencies using ‘-MajorMinor 1’, base counts for each site with ‘-doCounts 1’, ran the ‘-doMAF 1’ flag to determine minor allele frequency, and found posterior probabilities with the ‘-doPost 1’ flag. We filtered the geno file with the *‘*MAF’ flag to remove sites with a minor allele frequency less than 0.01. We computed a correlation matrix from the geno file with ngsPopGen^72,74^ using the *ngsCovar* command. The output covariate file was input into R^71^ where we used the *prcomp* function from ggfortify^75,76^ to perform PCA. Based on evidence of clustering by library preparation method in our initial PCA results (see Supporting Information 1; Figure S1), we repeated the analysis separately for historical samples and modern samples (Figure 2 A & B). We were particularly interested in whether there was genetic evidence for distinct subpopulations and whether patterns of genetic differentiation correlated with geographic distance within each analysis. We note that we also performed several analyses to identify relatives but did not find evidence of closely related individuals in our dataset (Supporting Information 2; Figure S2).

**Figure 2.**
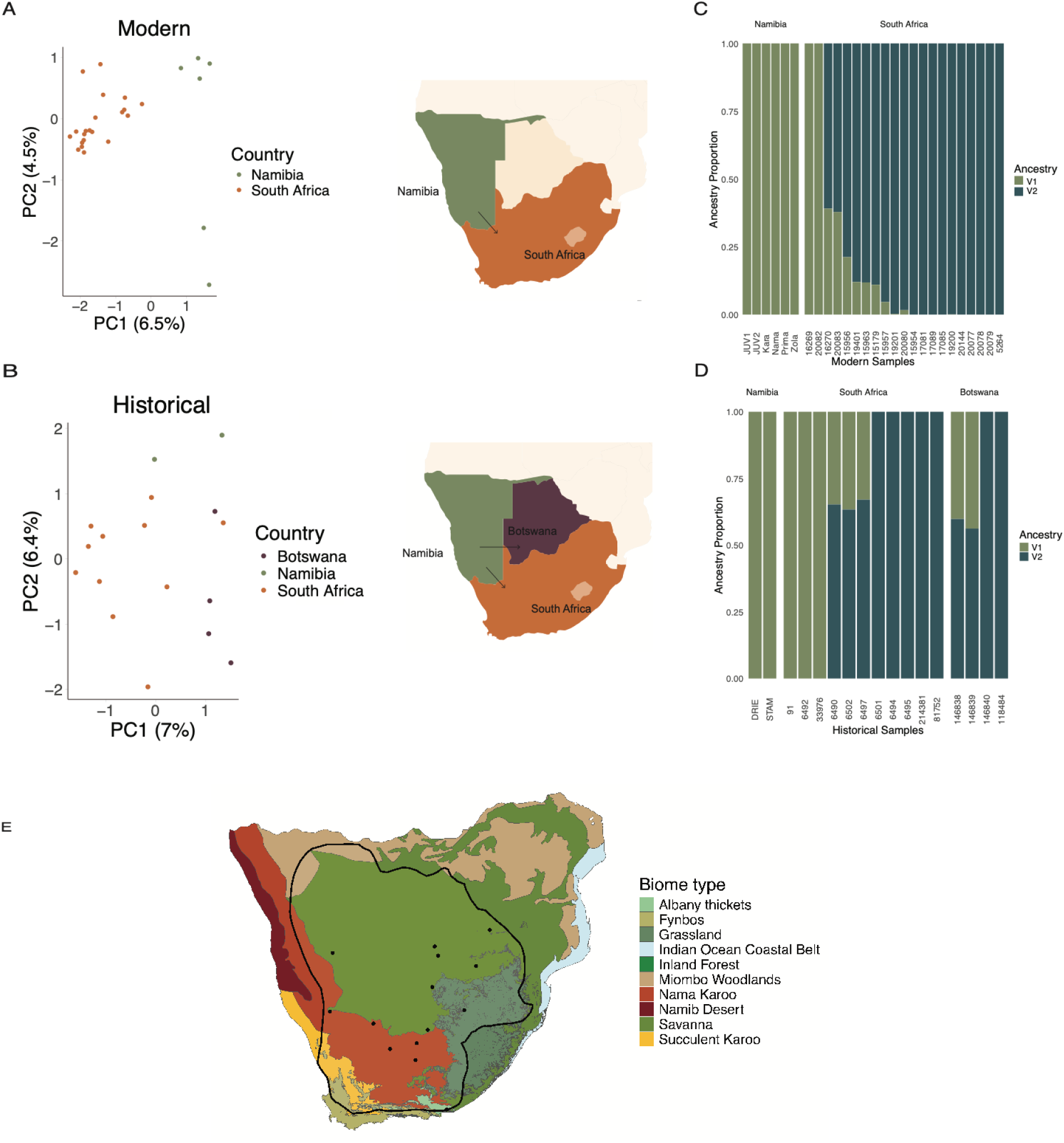
Population structure across the species range in black-footed cats. **(A-B)** Principal component analysis assessing population structure in **(A)** modern and **(B)** historic black-footed cat samples across the species distribution (PC1 and PC2 are plotted here). Samples are colored by country of origin (Botswana – eggplant, Namibia – olive, and South Africa – orange**). (C)** Modern and **(D)** historic NGSadmix plots for analysis of the data with K=2. Colors represent the two inferred ancestry groups. Samples are separated by country of origin which is labeled in the subheader and approximately ordered by increasing distance from Namibian sites. To the left of each admixture plot, there is a map of southern Africa. The arrows in the map show the direction in which samples were ordered by distance. (**E**) Map of biomes covering southern Africa. Different biomes are represented with different colors. Legend lists biome names.

### Ancestry proportions

To contextualize our PCA results, we aimed to investigate whether admixture between populations could be influencing observed population structure. In initial admixture analyses, we attempted to run ADMIXTURE^77^ to estimate ancestry proportions (Supporting Information 3; Figure S3). We found in this analysis resulted in unexpected associations with library preparation methods, likely driven by differences in sequence quality between historic and modern samples (Supporting Information 3). As an alternative approach to assess admixture between individuals, we analyzed our samples using the program NGSadmix^78^. NGSadmix estimates individual admixture proportions using genotype likelihoods instead of genotype calls to reduce reference bias and related errors in low coverage data. We downsampled data in BAM format from all individuals to ∼1X to avoid artifacts driven by coverage differences between individuals (see Methods above). We generated a BEAGLE input file using the ANGSD^73^ flag ‘-doGlf 2’ using the downsampled 1X BAM files before proceeding to run NGSadmixwith default parameters for multiple cluster options (K=2,4,5,10). NGSadmix estimated K=2 as the most likely number of ancestral populations so we continued with this value to visualize results using ggplot2^70^ and reshaper2 (Figure 2 C & D)^79^. Results from NGSadmix were suggestive of gene flow between geographically distinct populations, raising the possibility that individuals are moving across their range. Due to the black-footed cat’s known habitat preferences, we investigated biome variation across the species distribution. To visualize biomes concurrently with the species distribution, we plotted south African biome data^80^ and overlayed the species distribution using R (Figure 3).

**Figure 3.**
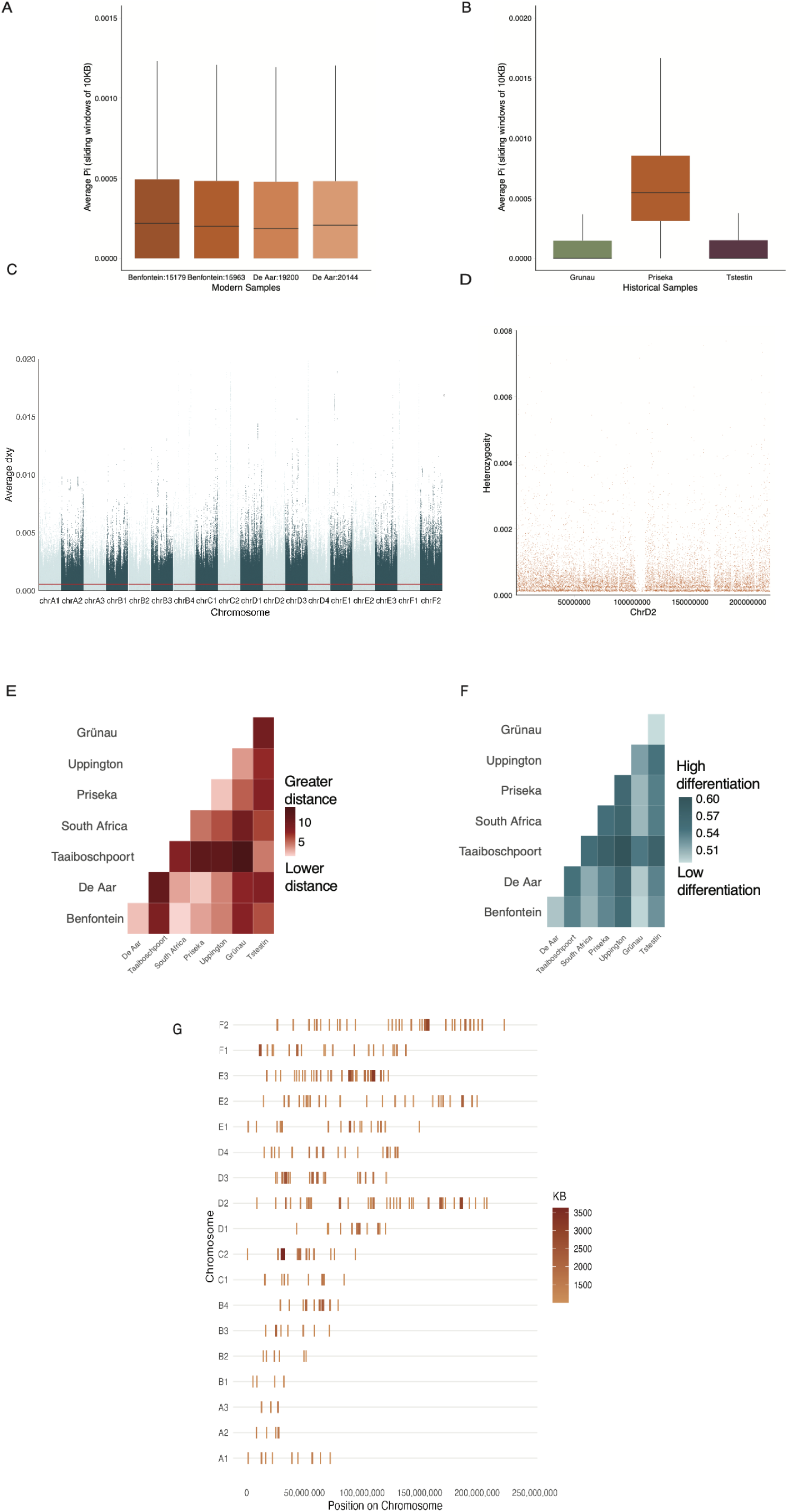
Genetic diversity and differentiation across the genome of black-footed cats. **(A)** Distribution of pairwise nucleotide diversity, *π*, in modern individuals (N=4) with at least 15X coverage; all of these individuals were sampled from two sites in South Africa. **B)** Genome wide average *π* in 10 KB sliding windows for historic individuals, all downsampled to 7X coverage, across the three countries where black-footed cats occur (Grünau - Namibia; Prieska - South Africa; Tstestin - Botswana). **(C)** Average pairwise sequence divergence (d_XY_) across the genome for South African individuals compared to Namibian individuals with coverage ≥5X. Only modern samples were used in this analysis**. (D)** Zoomed in view of average π in 10 kb windows in one individual across chromosome D2 spanning from 165 Mbp to 180 Mbp, a region inferred to fall in an ROH tract. **(E)** Heatmap of locality distance between sites shown in **F**. Darker color represents pairs of sites with greater geographic distance. **(F)** Comparisons of d_XY_ between samples collected from different localities, limiting to individuals with greater than 5X coverage. Sequence divergence (% d_XY_ per basepair) between localities is represented using blue color scale. Darker coloration represents higher genetic divergence. **(G)** Positions of detected runs of homozygosity (ROH) across the 19 black-footed cat chromosomes from one modern, South African individual. Only tracts larger than 1000 KB are shown. Orange color gradient shows KB size scale with darker shades corresponding to large ROH sizes.

### Analyzing genetic diversity and pairwise sequence divergence with Pixy

We calculated population genetic summary statistics, including nucleotide diversity (*π*) and pairwise sequence divergence (d_XY_), for each sample with sufficient coverage using the program Pixy^81^ (Table S1). Pixy accounts for uncertainty in the number of invariant sites and accounts for missing data in both the denominator and numerator in final calculations. Because individuals with very low coverage will have allelic dropout at heterozygous sites, we aimed for 10X coverage of samples included in this analysis. To measure nucleotide diversity within these individuals across the genome, we calculated the proportion of heterozygous sites within each 10 kb windows (Figure 3A). Due to potential technical bias arising from differences in sequence quality, we repeated this analysis in the high coverage preserved samples separately (Figure 3B). For calculations of d_XY_, Pixy^81^ calculates average pairwise sequence differences between individuals from different populations in a user-specified 10 kb sliding windows across the genome. While we had aimed for 10X coverage of these samples, in practice some individuals did not reach this threshold, so we ultimately repeated our analyses downsampling individuals to 7X coverage to eliminate coverage differences between these individuals (Figure 3B; 3E).

### Runs of homozygosity

We used PLINK^82,83^ v. 2.0a7 to estimate runs of homozygosity (ROH) in high coverage non-preserved samples. PLINK requires PED files for input so we converted our VCF containing 15X coverage individuals to a BED file using the *make-bed* command. PLINK is configured for the human genome by default so we included *allow-extra-chr* command to accept non-human chromosome codes and excluded sex chromosomes with the *no-sex*. We converted the BED file to a PED file in plink using the ‘recode’ flag. We then used our PED file to estimate ROH using the *–homozyg* command. We used the default ROH parameters in plink except that we modified *homozyg-window-missing* and *homozyg-window-het* to allow a window to contain 10 missing calls and 2 heterozygous calls. We also explored different values for these parameters and found our results to be relatively insensitive to the exact values used.

### Estimating historical and recent effective population sizes

We performed demographic analysis using three samples from geographically distinct collection sites (Botswana, Namibia, and South Africa; Figure 4). These samples were targeted for higher coverage data collection and underwent the same library preparation and data processing protocols. Based on initial analysis (Supporting Information 4; Figure S4), we suspected that variation in coverage among these samples was impacting our results, so we down-sampled data to 7X coverage. For each individual BAM file, we generated consensus sequences in FASTQ format using SAMtools^41^ and BCFtools^42^ following the methods described by Palkopoulou et al.^84^. Briefly, we indexed BAM files with BCFtools^42^ then combined *mpileup* and *vcf2fq* commands from vcftuils.pl^64^ to generate a consensus sequence for each individual and chromosome while filtering for a minimum base quality of 30, mapping quality of 30, and RMS mapping quality of 30. The consensus FASTQ^56^ files were used for pairwise sequentially Markovian coalescent^85^ (PSMC) which was run under default parameters (64 time intervals: 4+25*2+4+6). We performed bootstrapping by subsampling the consensus sequence with replacement (50 kb) and ran PSMC^85^ 100 times for bootstrapped replicates of each sample. For visualization, we scaled the data by a mutation rate of 0.86 x10^-8^ per bp per generation based on available estimates from the domestic cat^86^. There is some uncertainty of the black-footed cat generation time, but we use a generation time of 3 years in this study. We then plotted results for each individual using the PSMC perl script psmc_plot.pl^39^.

**Figure 4.**
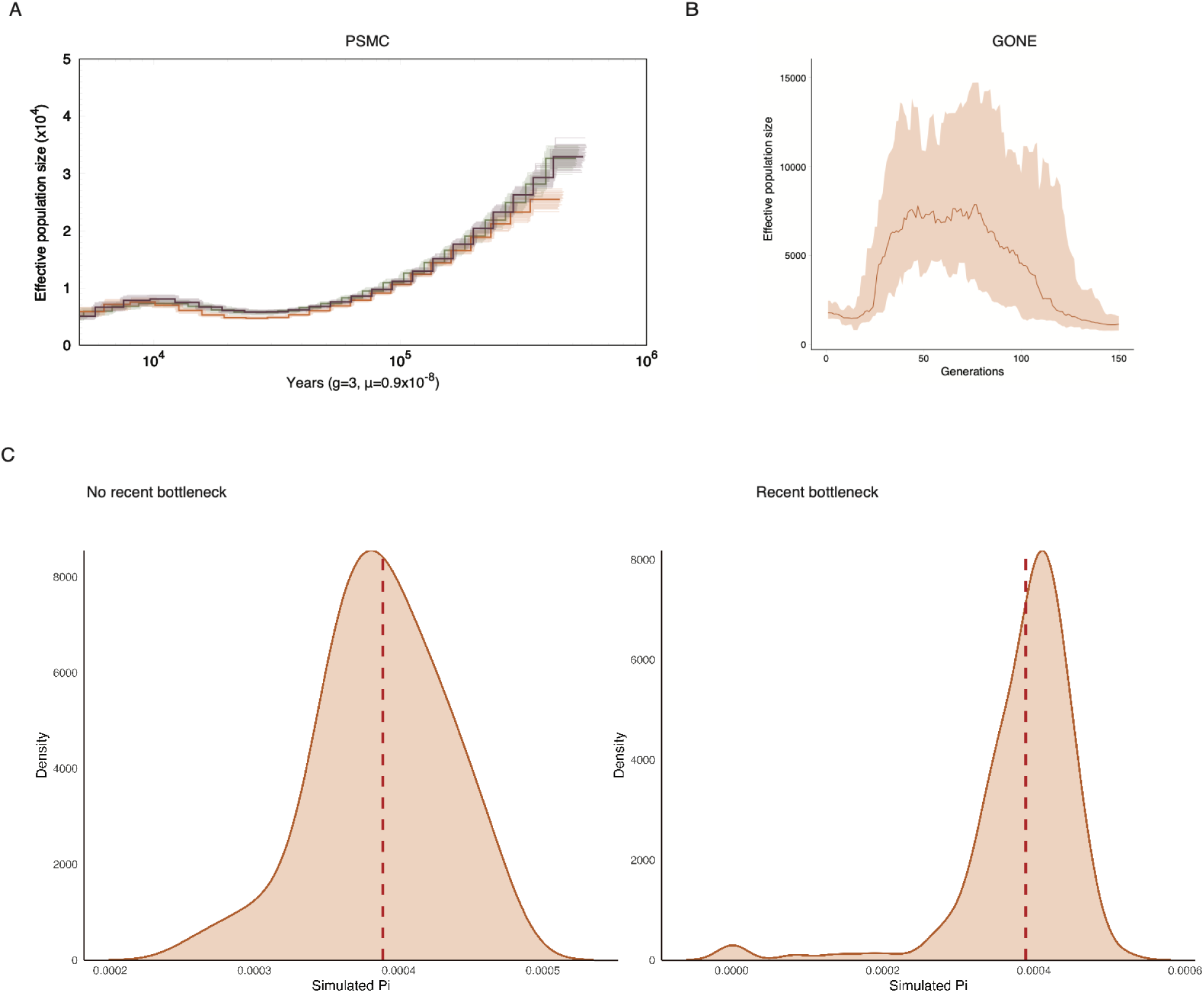
Demographic history. **(A)** Effective population size (N_e_) estimates were reconstructed using the Pairwise Sequential Markovian Coalescent^85^. We performed this analysis for three high-coverage black-footed cats sampled from three geographical regions (Botswana – eggplant, Namibia – olive, and South Africa – orange). For visualization, we assumed a mutation rate of 0.86 x 10^-8^ per basepair per generation and a generation time of three years^86^. Lighter lines show the results of 100 bootstrap replicates for each sample. (**B)** GONE estimates of demographic history within the past 150 generations from high coverage (>10X) South African individuals (from median estimates are shown by dark orange line). Effective population size remains modest but suggests population decline within the last 50 generations. 95% confidence intervals were produced from 40 replicates and are shown in orange shading. **(C)** SLiM simulation results from 150 replicate simulations of two different demographic scenarios. Distributions show expected genetic diversity under scenarios of a constant historical population size of 11,500 individuals (left) and a constant size of 11,500 individuals followed by a recent contraction to 2,800 to model GONE results (right). The red dashed line in each distribution corresponds to the observed *π* in the empirical data (0.00039 per bp).

Our results indicated that the geographically distinct samples had a very similar population history (Figure 4A). Thus, we focused on the South African sample for further analyses and simulations. Using this South African sample, we continued to investigate historical population trends and explore how these trends could result in present-day *π*. We calculated the weighted time-averaged population size based on our PSMC^85^ results from this individual between the last 2,000-100,000 years, excluding time points where we lose resolution in the recent and distant past. This resulted in an estimate of 1.15 x 10^4^ for the weighted historical population size of the black-footed cat. We next used simulations implemented in SLiM^87,88^ to estimate the expected *π* for a population with this historical size, a mutation rate of 0.86 x10^-8^ per bp per generation, and recombination rate of 1.74 x10^-8^ per bp per generation (both inferred from the domestic cat; Figure 4C)^86^. We performed simulations using a burn-in of 10 N_e_ generations, followed by a simulation period of 100,000 generations, with a constant simulated population size of 1.15 x 10^4^. We simulated 1 Mb of sequence and used the output VCF function in SLiM^87,88^ to output data for one individual. From this VCF we calculated *π* in the simulation and compared this value to the observed *π* in our data. We repeated this procedure 150 times.

To investigate demographic trends within the past 150 generations, we used GONE^89^ to estimate effective population size from linkage disequilibrium patterns and recombination rate (Figure 4B)^90^. In contrast to PSMC^85^ which has lower resolution in the recent past, GONE^89^ can be applied to estimate recent effective population size. We generated MAP and PED files from the VCF containing South African samples with ≥ 15X coverage in VCFtools^64^ using the ‘plink’ flag with a minimum allele frequency filter ‘-minMAF’ of 0.01. We ran GONE^89^ under default parameters with 40 replicates, which were used to estimate confidence intervals, for inferred effective population size then visualized results in R using ggplot2^70^ with a script adapted from Kardos et al. (2023)^91^ and de Greef et al. (2024)^92^. We initially ran GONE^89^ using the domestic cat (*Felis catus*) genome-wide sex-averaged recombination rate of 1.9 cM/Mb since recombination rate estimates are not available for the black-footed cat^93^. We found that GONE^89^ was highly sensitive to the user-provided recombination rate, so we repeated analyses exploring multiple values for this parameter (Supporting Information 5; Figure S5).

Based on evidence of a recent population bottleneck from the GONE analysis, we repeated SLiM simulations as described above, except that we implemented a bottleneck to 2,800 diploid individuals at 150 generations before the present. We evaluated the effect that a recent bottleneck would have on *π* based on these simulation results.

## Results

### Genome Assembly

Our initial PacBio assembly with hifiasm^35,37^ resulted in a primary assembly totaling 2.6 Gb across 5,115 contigs, with a Contig N50 of ∼1.5 Mb (Table S1). Subsequent Hi-C data incorporation yielded a primary assembly of 2,456 scaffolds, with a Scaffold N50 of 142.6 Mb (Figure S7). This new assembly reduces the total number of scaffolds by 10-fold compared to the previously available DNAZoo draft assembly. More than 94% of Carnivora single gene BUSCO orthologs are present and complete compared to 90% in the assembly by DNAZoo^48,94^. Another black-footed cat genome assembly recently became available^28^, generated using PacBio data and used the same publicly available Hi-C reads. While this assembly has greater contiguity (Scaffold N50; 146.9Mb, Contig N50; 37.19 Mb), BUSCO^46,48^ scores were similar (97%), so we opted to continue with the assembly we had generated.

Past work has indicated that Felidae tend to have high synteny, even across divergent species^95^. We thus used the chromosome assignments from Felcat9 in conjunction with the visualization program Circa (http://omgenomics.com/circa) to assign chromosome IDs to the scaffolds in our assembly (Figure 1D)^96,97^. The assembly we generated contains all 19 chromosomes including the X chromosome (matching those in the Felcat9 assembly)^94^. MUMmer^50^ results showed high similarity between chromosomes in the two assemblies (Figure 1E)^28^.

### Population Structure

We found that modern individuals separated along PC1 roughly by country of origin (Figure 2A). Little variance is explained by PC2 with modern Namibia samples clustering tightly together. We noticed the presence of three outlier individuals among the modern South Africa individuals. We examined these individuals for potential technical issues that could drive them to be outliers but did not find evidence of problems with these samples (Supporting Information 1). These samples did not have precise location data and likely represent samples collected further from the Benfontein site (e.g. opportunistically sampled from vehicular collisions; Alexander Sliwa, personal observation). The PCA of historical individuals exhibited weaker clustering by country of origin (Figure 2B) but it is unclear if reduced signals of geographic differentiation are attributable to lower overall data quality obscuring the signal of interest or a true difference in population structure over time.

### Recent Admixture

For both the modern and historic NGSadmix^78^ analyses, we identified K=2 as having the lowest cross validation value. For K=2, NGSadmix results for the modern samples supported evidence of one ancestry group present in Namibian samples (Figure 2C). Unlike the Namibian samples, the South African individuals were inferred to contain two ancestry groups. Analysis of historical data included samples from all three countries in the black-footed cat range (South Africa, Namibia, and Botswana). We found a similar trend as in our analysis of modern individuals. Historic Namibian individuals were again inferred to represent one ancestry group (Figure 2D) while Botswanan and South African individuals were inferred to represent two ancestry groups. These results add to our inferences from population structure and d_xy_ results suggesting that the species may have higher migration between geographical locations than previously thought.

### Quantifying present day genetic variation

We estimated that *π* was ∼0.0004 per basepair, consistent with previous estimates from zoo and museum specimens (Figure 3 A & B)^28^. Analysis of pairwise d_XY_ revealed little genetic differentiation between sampling localities. This is surprising given observational studies that have suggested that dispersal between populations is limited^19^. To further investigate these patterns, we analyzed the correlation between pairwise collection site distance and pairwise d_XY_. Reassuringly, we found that samples that were furthest from each other geographically had higher d_XY_, consistent with a model of isolation by distance, although this relationship was not statistically significant (R=0.3, p=0.07; Figure E & F). However, in general, observed d_XY_ between sampling sites was only slightly higher than within population diversity (d_XY_ ∼ 0.00065; Figure 3C). We also evaluated local variation in d_XY_ along the chromosome. Genome-wide we observed that d_XY_ was typically low, interrupted with some peaks of high differentiation throughout the genome. These peaks could be attributed to selection against gene flow or genomic regions with lower local recombination rates and higher background selection (Figure 3D). Together, this suggests higher than anticipated connectivity and low genetic differentiation between black-footed cat populations across the species range.

Low standing genetic diversity is often associated with inbreeding. However, we found very few long (i.e. 3 Mb or greater) ROH present in the sampled black-footed cat genomes (Figure 3G; ROH size class noted in Figure S6). We tested different parameter sets to investigate whether low genetic diversity combined with high rates of error might be disrupting the detection of ROH (see Supporting Information 6) but continued to find little evidence of long ROH. This suggests that despite low genetic diversity, there is not clear evidence for recent inbreeding in our dataset. These results are concordant with visualizations of the raw data (Figure 3D). These visualizations highlight some areas of reduced genetic diversity compared to the background, but these regions tend to be shorter than 3 Mb (Figure S6).

### Demographic history

We reconstructed the demographic history of high-coverage individuals representing each geographic group (Botswana, Namibia, and South Africa) using PSMC^85^. Bootstrap resampling of the data suggested that we lose resolution to infer population size in the recent past, ∼1,000 years before the present. We found that population histories from different sampling locations mirror each other. PSMC^85^ analyses suggest that populations declined to around 8,000 individuals by ∼10 kya before the present, then increased to 10,000 individuals over a ∼5 kya period, possibly coinciding with the end of the last glacial period when drying events altered vegetation and produced more arid biomes (Figure 4A)^98^. Further declines in effective population size were inferred after this period. We note, however, that uncertainty in the mutation rate and generation time could impact the accuracy of these estimates and concordance with climatic events. Integrating over periods of increases and declines in population size, the weighted average historical effective population size for these species was inferred to be ∼11,500.

Since PSMC^85^ only reliably estimates population size in the more distant past, we used GONE^89^ as an alternative method to estimate recent effective population size. GONE^89^ results suggested that black-footed cats had a population size of ∼8,000 around 100 generations before the present, followed by substantial decline in the last 50 generations (Figure 4B) to an estimated size of ∼2,800 individuals. GONE^89^ population size estimates are dependent on user input recombination rate to quantify linkage disequilibrium and use these patterns to infer recent population size^90^. Since no recombination map is available for the black-footed cat, we used estimates from the domestic cat and varied this parameter to test the program’s sensitivity to misspecification^93^. We found that GONE^89^ was highly sensitive to the recombination rate parameter and thus these results should be interpreted with caution. We discuss these issues in more detail in Supporting Information 5 and Figure S5.

To further investigate these possible demographic events, we implemented simulations in the forward-time population simulator SLiM^87,88^. We were interested in determining whether inferred population histories from PSMC and from GONE were consistent with our empirical data. To do so, we first implemented simulations in SLiM using the likely black-footed cat mutation rate, recombination rate, and the inferred time averaged population size of ∼11,500 from the PSMC analysis. Strikingly, we found close concordance between the simulated genetic diversity in these SLiM simulations and the observed levels of genetic diversity in our data, with an average genetic diversity in simulations of π = 0.00039 (Figure 4C; 95% confidence intervals: 0.00028-0.00047). This suggests that unlike other felids, the levels of genetic diversity observed in black-footed cats can be explained by their modest historical population size, without needing to invoke the presence of a recent bottleneck^99^. However, motivated by the GONE results, we repeated these simulations implementing a strong bottleneck to ∼2,800 individuals in the past 150 generations. We found this recent reduction in population size was also concordant with observed levels of genetic diversity in our data, suggesting that a recent bottleneck of this magnitude would not be severe enough to have a strong impact on genetic diversity (average simulated π = 0.00038; Figure 4C; 95% confidence intervals: 0.00017-0.00047).

## Discussion

Black-footed cats are a charismatic species whose population genetic history and contemporary structure are remarkably understudied. Despite their description in 1824, black-footed cats were largely overlooked until the establishment of the BFCWG in 1992. Observational data collected by this group suggested that the species has experienced population decline in the past 20 years, potentially driven by human disturbance^19^. These studies reported impacts of human disturbance, including mortality due to trapping, black-footed cats consuming poisons meant for larger predators, and habitat degradation, as potential factors contributing to decline^22^. Together, these findings led the IUCN Red List to categorize the species as vulnerable to extinction in 2002. However, in 2015 during their re-assessment, the IUCN underscored important data deficiencies for this species, and lack of information from across its range^19^. Here, we aimed to significantly improve our understanding of genetic diversity and demographic history in black-footed cats using genomic sequencing to inform their current extinction risk. This represents the first genomic study to investigate ecological and evolutionary dynamics in natural populations for this species.

In this study, we assembled a high-quality reference genome and collected genome-wide data for 44 free-roaming cats distributed across the species range using modern and historic specimens (collected between 1818-2009). We found weak evidence for population structure based on PCA results, which could indicate that black-footed cat populations are more connected than previously predicted. Black-footed cats are known to inhabit home ranges averaging in size from 10 km^2^ (females) to 20.7 km^2^ (males) depending on food availability, and previous work predicted the species has maximum dispersal distance of 50 km^19,21,100^. However, we find very weak genetic differentiation between populations, only slightly exceeding within population diversity, which could point to unexpected connectivity between different populations of black-footed cats. Alternatively, low genetic differentiation could be explained by the present-day populations having recently arisen from a common historical source population.

Our admixture results provide further support for some connectivity between black-footed cat populations. We infer the presence of two ancestry groups in our dataset, with evidence for admixture between the Namibian populations and populations in South Africa and Botswana. Southern Africa is rich in biome diversity with extremely distinct environments across this geographical landscape, ranging from deserts to fynbos. These biomes vary in climate and vegetation which affects species composition, suitable habitat, and food sources for black-footed cats. Black-footed cats are thought to prefer arid to semi-arid environments, with the majority of their species distribution ranging over the Nama Karoo and savanna ecosystems (Figure 2E). Thus, a corridor of suitable habitat may have facilitated movement of individuals between Namibia and eastern locations, influencing the admixture patterns we find in this study. These types of ecological drivers can strongly influence genetic diversity within, and genetic differentiation between, populations ^101,102^. Consistent with a history of genetic exchange between populations, analysis of pairwise sequence divergence (d_xy_) suggested that despite some evidence of isolation by distance, genetic differentiation between individuals originating from different localities was low.

Given our inferences about population structure and gene flow, we predicted black-footed cats would have relatively higher genetic diversity compared to other felids who are at risk of extinction. Surprisingly, our findings suggest that black-footed cats exhibit remarkably low genetic diversity, with estimates even lower than other iconic threatened cat species like the African lion (*Panthera leo*)^103^ and tiger (*P. tigris*)^99^. Despite low diversity, individuals do not exhibit signs of recent inbreeding, with few observed ROH tracts longer than 3 Mb. While we were unable to infer ROH in historical samples due to data quality issues, analyses of ROH in the earliest available samples from the 1800s would be a valuable contribution if future studies are able to collect higher coverage data from such samples.

Analyses of historical population sizes using PSMC suggest that black-footed cats historically had a moderate effective population size compared to large felids, with a time-averaged population size estimate of ∼11,500 individuals. This inferred historical population size differs substantially from other felids such as lions and leopards (*P. pardus*) which have estimated historical populations sizes ranging from 50,000-150,000^99^. Even when we repeated our analysis using the same mutation rate applied in these studies, the estimated effective population size for black-footed cats remained smaller compared to the lion and leopard (Figure S4B-C). Simulations suggest the low levels of genetic diversity observed in present day black-footed cat populations are consistent with the demographic history inferred from PSMC and need not invoke recent declines in population size. However, analyses using GONE^89^, which estimates population size in the recent past, were consistent with a bottleneck in the last 50 generations (Figure 4B). While these findings are concordant with the results of monitoring efforts by the BFCWG, we also found that our analysis was very sensitive to small changes in the user-provided recombination rate (Supporting Information 5; Figure S5). Thus, we interpret these results cautiously but note that the estimated contraction coincides with in farming practices in the region, notably the transition to commercial farming. During this period, black-footed cat populations in South Africa may have experienced much higher livestock numbers and resulting habitat degradation. Farming practices introduced during colonial times often devastate local wildlife through mass poisoning, overgrazing, and loss of apex predators or migratory herds^104–107^. These practices continue into present day and may be continuing to degrade black-footed cat habitats.

The extinction vortex is a hypothesis used to describe the genetic mechanisms contributing to extinction via a four-component negative feedback cycle^4^. Ecological and observational studies have indicated that black-footed cats are experiencing increasing habitat fragmentation and appear to be declining at key study sites, demographic trends that we may be detecting in GONE analyses. However, we see little evidence of inbreeding in our analyses and speculate that higher dispersal or connectivity between populations may partially counteract the impacts of possible population size reductions. Moreover, our analyses suggest that black-footed cats have long persisted at more modest population sizes compared to other felids, which has likely reshaped the landscape of recessive deleterious alleles segregating in this species and the strength of inbreeding depression. These complex relationships between genetic diversity, population resilience, and genetic load are consistent with emerging genomic studies in other natural populations of conservation concern, highlighting a need to revisit classic hypotheses to better understand the impacts of demographic fluctuations, gene flow, and genetic variation on population resilience^108^.

Here, we used whole genome sequencing of free ranging individuals across the black-footed cat range to infer population structure, connectivity, and demographic history. We find that black-footed cats have markedly lower historical effective population sizes compared to other wild felids, contributing to low standing genetic diversity. Despite this, black-footed cat populations do not show signs of recent inbreeding and appear to have higher than expected connectivity across geographically diverse populations. Given this surprising finding, human development may pose a threat to this species in future years. Research has already shown that anthropogenic factors, such as farming practices, contribute to black-footed cat deaths in the wild^19^. As southern Africa continues to develop, with increased roads and expanded farmlands, these small cats may experience difficulty maintaining population connectivity, which could result in increased inbreeding. Further exploration of black-footed cats could improve our understanding of how metrics like gene flow and migration can be leveraged to support conservation efforts. Integrating genomic data into IUCN Red List assessments will provide increased support for species assessments and illuminate potential paths towards conversation management of this important, yet poorly known, species.

## Acknowledgements

We would like to acknowledge the efforts of the Black-footed Cat Working Group and the Black-footed Cat Research Project Namibia who contributed samples, expertise and time to this study. In addition, we take this opportunity to thank the San Diego Zoo Wildlife Alliance for assistance procuring international samples and sample storage prior to the project. We also thank the McGregor Museum (Kimberley, South Africa), the American Museum of Natural History (New York City, NY), Museum of Vertebrate Zoology (University of California- Berkeley), and the Harvard Museum of Comparative Zoology for providing subsamples from their museum specimens. We thank Bill Murphy and Gang Li for helpful discussion and permission to reproduce past phylogenetic analyses. This material is based upon work supported by the National Science Foundation Graduate Research Fellowship under grant No. DGE-1656518 to VBG, funding from National Institute of Health (NIH) Cellular, Molecular Biology Training Grant (NIH 5 T32 GM007276) to VBG, and funding to MS through the Freeman Hrabowski Scholars program.

