## Supplemental text for "Genetic Diversity and Population Structure of the Black-Footed Cat: Insights into *Felis*’s Deadliest Predator"

[10]Department of Evolution Ecology & Organismal Biology. University of California, Riverside

[11]Howard Hughes Medical Institute, Stanford, CA

### Supporting Information

*Supporting Information 1. Correcting for technical biases in the principal component analysis*

We performed principal components analysis (PCA) on our data to investigate genetic differentiation between individuals and possible evidence of geographic structure. Initially, we included all samples, regardless of data quality or source, in the same analysis. This dataset ranged in coverage from ~1-30X (n=44) and included both modern and historical individuals. Since our data included low coverage individuals, we used ANGSD v. 0.931[1,2] to generate genotype likelihoods at each SNP for each individual. We performed PCA analysis on BAM files using the same pipeline for analysis and visualization from the main text (see Methods).

When examining these initial results, we found that samples clearly separated by modern and historical individuals along PC1, suggesting that observed results were dominated by the differences in coverage, library preparation method, and/or sample quality (Figure S1A). We also found significant correlations between PC loadings and coverage in this initial analysis (PC1, R= -0.61, p=0.0000129; PC2, R=0.14, p=0.36). To address this coverage bias, we first re-ran the PCA separately on high (≥10X) and low (<10X) coverage individuals. Although this approach alleviated issues related to the correlation between coverage and PC loading (PC1, R=0.21, p=0.29; PC2, R=0.22, p=0.30), a strong separation on PC1 between the historic and modern samples in the high coverage analysis remained (Figure S1B). We interpret this observed pattern as evidence for technical bias caused by differences in sequence quality (Figure S1C). We also detected continued correlation between coverage and PC2 loadings in this stratified analysis (PC1, R=-0.45, p=0.06; PC2, R= -0.68, p=0.001271). In an attempt to account for these layered technical issues and improve biological interpretation, we repeated this analysis while separating samples by library preparation method (non-preserved (all samples using the non-preserved library prep were South African) or preserved, see main text Methods; Figure S1D-E) and with BAM files down-sampled to ~1X coverage to eliminate bias caused by differences in coverage (Figure S1F). We performed the PCA across three separate analyses: one with all individuals, and then two analyses separately for the historic and modern samples. The PCA plots generated from this analysis (see Methods in main text) showed clustering by sample origin in both historic and modern PCA analyses (main text; Figure 2A-B).

*Supporting Information 2. Relatedness analysis*

Samples collected for this study were derived from free-roaming wild-caught individuals and we did not have access to any pedigree information on potential relatedness between individuals. We aimed to explore population structure within black-footed cats to understand the intraspecies interactions, but related individuals could impact population structure analyses since they violate the assumption that no relatives are present in the dataset. We estimated the relatedness of all individuals in our VCF file (n=44) using the VCFtools[3] '--relatedness2' flag, based on the KING method of Manichaikul et al.[4] and then visualized results using ggplot2[5].

We found that none of the individuals in our dataset show evidence of close relatedness to other individuals analyzed (Figure S2). The relatedness metric from KING compares all individuals in the VCF to each other. Relatedness estimates close to zero indicate that individuals are unlikely to be related, while higher estimates are consistent with a higher probability of relatedness. Across all pairs of individuals, the estimated relatedness statistic was below 0.25, suggesting that our dataset did not include close relatives. The self-comparison metric (i.e. comparing individuals to themselves) showed relatedness of 0.5 meeting expectations for a high degree of similarity in self-comparison.

*Supporting Information 3. Admixture*

In our PCA results (main text, Figure 2A & B), we found less distinct clustering than we anticipated considering that geographic distances between samples were large relative to the estimated dispersal distances for the species. To investigate potential evidence of recent admixture, we first analyzed our samples using the program ADMIXTURE[6]. We used our VCF to generate a PLINK BED file format using PLINK v. 2.0a2[7] and the '--make-bed' with MAF threshold of 0.02. ADMIXTURE was designed for human datasets and anticipates human chromosome names in order to identify each chromosome during cross validation. To accommodate non-human chromosome names, we replaced black-footed cat chromosomes names with numbers in our BED input file before running the cross validation '--cv' loop. We ran cross validation within ADMIXTURE for K=2, K=3, K=5, K=10. We identified the K=2 cluster as having the lowest cross validation value and proceeded with K=2 for analysis of our dataset. We visualized the results in R[8] through ggplot2[5].

In these initial ADMIXTURE results, we found that the detected "ancestry clusters" largely corresponded to sampling date and library type, suggesting differences in sample quality were influencing findings (Figure S3A). We attempted to re-run ADMIXTURE with only modern samples to avoid artifacts generated from quality differences between historic and modern samples (Figure S3B). In this analysis, we identified K=2 as the best cross validation value and proceeded with this value for analysis and visualization. ADMIXTURE results for the modern samples found one ancestry group present in Namibian samples but a mix of two ancestry groups in South African individuals. These results could be consistent with higher than expected migration between geographical locations, with Namibian migrants contributing to the South African populations. However, we were interested in analyzing historical samples as well, while accounting for issues that could arise due to lower sequence quality. To correct for the difference in sequence quality, we ran NGSadmix[9] (see main text methods) on the historic and modern samples separately. The NGSadmix analysis also resulted the identification of K=2 as the best cross validation value for both historical and modern samples (main text; Figure 2C-D).

*Supporting Information 4. Pairwise Sequential Markovian Coalescent*

We conducted demographic analysis using three samples, representing one high coverage individual from each of the three countries within the dataset (Botswana, Namibia, and South Africa). We generated consensus sequences in FASTQ format from each individual BAM file using SAMtools[10] and BCFtools[11] as described by Palkopoulou et al[12] and ran PSMC[13] as described in the main text (see Methods). The results of this initial analysis suggested similar trends in demographic history across geographic locations (Figure S4A). However, the results for the Namibian individual, which by chance had lower coverage than the other samples (~7x vs ~10x), appeared to be shifted along the x-axis. While distinct population history for this location is possible, we suspected that this shift along the x-axis could also be driven by differences in coverage. To investigate this, we down sampled the South Africa and Botswana representative samples to the same coverage (~7X) to produce the PSMC results for each sample presented in the main text (Figure 4A).

Additionally, we made comparisons of the inferred demographic history of the black-footed cat demographic history to the African lion (*Panthera leo*) and the African leopard (*Panthera pardus*). For this comparison, we input our PSMC results into psmc_plot.pl and applied a mutation rate of $5.4 \times 10^{-9}$ and $7.6 \times 10^{-9}$ for the lion and leopard respectively. We found that using either the lion or leopard mutation rate did not substantial change our results (Figure S4B & C), supporting the conclusion that black-footed cats have had modest historical population sizes when compared to large felids.

*Supporting Information 5. Additional details on GONE parameter specifications*

We used the program GONE[15] to infer demographic history in the last 100 generations. Unlike PSMC [13] which loses resolution in the recent past, GONE is reported to accurately infer effective population size within the last 100 generations using patterns of linkage disequilibrium. Since estimates of average recombination rate are not available for black-footed cats, we tested the sensitivity of GONE results to changes in the recombination rate parameter by adjusting the cMMb parameter ± 10% (1.7 and 2.1 cM/Mb). We otherwise used default parameters in our analysis.

As discussed in the main text, using the domestic cat sex-averaged recombination rate of 1.9 cM/Mb, GONE results suggested that in the last 100 generations the peak effective population size was ~8,000 individuals and that around 50 generations ago, population size declined to below 2,800 individuals (main text; Figure 4B). However, we also found that results from GONE are extremely sensitive to the average recombination rate input by the user. When we modestly decreased the cMMb parameter from 1.9 to 1.7, we found a drastic increase in population size. The peak population size predicted in the last ~125 generations jumped from ~8,000 to ~30,000 with the confidence intervals extending to ~110,000 (Figure S5A). This inferred population size exceeds the peak historical population size inferred based on PSMC analysis. Given reports of black-footed cat population decline, this result is puzzling and is suggestive of high sensitivity of GONE to parameter misspecification. We continued to test parameter sensitivity by increasing the recombination rate from 1.9 to 2.1 which decreased the inferred population size (Figure S5B). For a recombination rate of 2.1, we found peak effective population size to be ~3,000 and a similar population size trajectory as observed in the run with cMMb set to 1.9. All GONE runs were performed with 40 replicates. We estimated confidence intervals for all runs of GONE[15] and found a wide range in population size estimates.

*Supporting Information 6. Alternative ROH analysis using GARLIC*

In the main text, we discuss evidence for runs of homozygosity (ROH) based on analysis of our data with PLINK[7]. As an additional approach, we quantified ROH with the program GARLIC[16]. Unlike PLINK[17] which uses an observation-based method that directly counts genotypes to infer ROH, GARLIC utilizes sliding windows with a model-based approach to infer significant ROH. Given the sensitivity of ROH analysis to genotyping errors, we only performed this analysis in high-coverage, modern samples. To generate GARLIC input, we used PLINK v. 2.0a2 with the "--allow-extra-chr", "--recode", and "--maf" set to 0.02 flags to convert the high coverage VCF to PLINK tfam and tped format. GARLIC was run using default parameters with assumed genotype error rate (--error) set to 0.01, the window size of SNPs used to calculate logarithm of odds score set to 100 (--winsize), and centromeres set at 0, since no centromere information is available black-footed cats.

The GARLIC analysis mirrored the results of PLINK overall, suggesting that the black-footed cat genomes contained tracts of small or medium ROH and very few long ROH tracts that would be indicative of recent inbreeding (Figure S6). However, we found that in the context of our data GARLIC[16] was very sensitive to the user-selected window size parameter, with results varying substantially between runs on the same individual with different window sizes. As a result, we use the PLINK analysis in the main text. However, we note that both analyses are consistent with little evidence for recent inbreeding (Figure 3G).

### Supplementary Figures

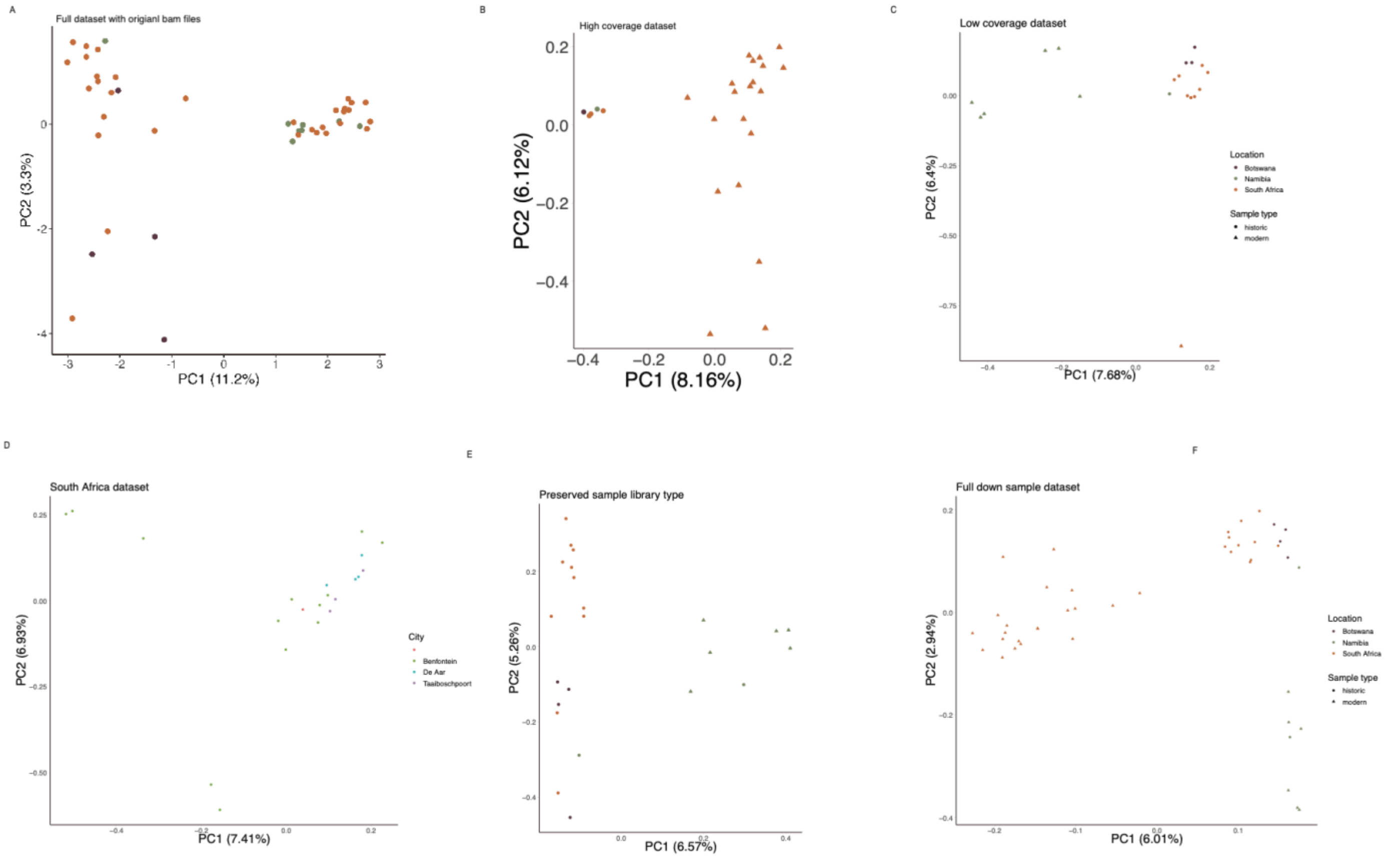


**Figure S1.** Principal component analysis generated with original BAM files containing the full dataset (comprised of historic and modern samples of a range of coverage). Population clustering is assessed in clustering along PC1 and PC2 using ANGSD. Samples are colored by country of origin (Botswana – eggplant, Namibia – olive, and South Africa – orange). Historical samples are marked by circle points and modern samples are marked by triangles. **A)** Results produced from individuals in the entire dataset without coverage normalization. Separation along PC1 corresponded with historical versus modern samples with coverage having a significant correlation with PC loadings. **B)** Analysis restricted to high coverage individuals of both historical and modern samples (≥10X). Historical and modern samples differentiated along PC1 with modern samples spreading along PC2. **C)** Analysis of low coverage individuals only (<10X). Historic and modern samples are noted by circles and triangles respectively. Historic and modern samples showed some separation along PC1, suggesting that technical differences were driving the signal, prompting us to separate these groups for further analysis. **D)** PCA of South African samples produced from non-preserved extraction and library prep. Most South African samples clustered together in PCA space regardless of precise sampling site within South Africa. Notably, we observe a substantial spread in placement in PCA space even among individuals collected from the black-footed cat long-term research site in Benfontein separated along PC1 and PC2. **E)** Analysis of data generated from the preserved library preparation method only, which included both historic samples and modern samples that needed to be prepared using this approach (e.g. due to low sample quality). Historic samples fell into two clusters on PC 1 and PC2 and separated from the modern samples. **F)** ANGSD PCA plot generated after downsampling all samples to 1X. Modern South Africa and Namibia samples separated along PC2 while historic samples segregated along PC2. These analyses prompted us to perform PCAs separately for historic and modern samples in the results presented in the main text.

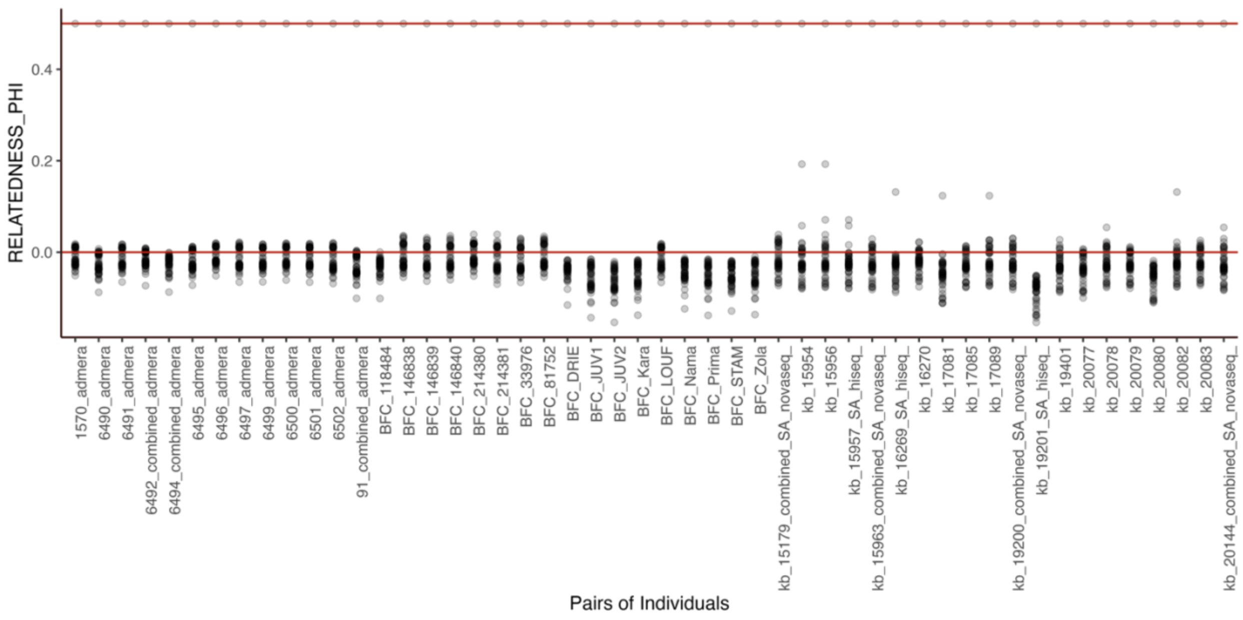


**Figure S2.** Estimated relatedness ϕ for individuals in our dataset based on the KING[4] method, implemented using VCFtools. The sample names are listed along the x-axis with the y-axis showing the estimated probability of finding identical alleles from random sampling an allele from each heterozygous individual. Every sample is compared to all the samples in the dataset including themselves. Each point represents a pair of samples being compared. Points close to 0 indicate pairs of samples that have a low probability of relatedness, whereas higher values indicate a higher probability of relatedness. The points at 0.5 for each sample are a positive control indicating the results of self-comparison.

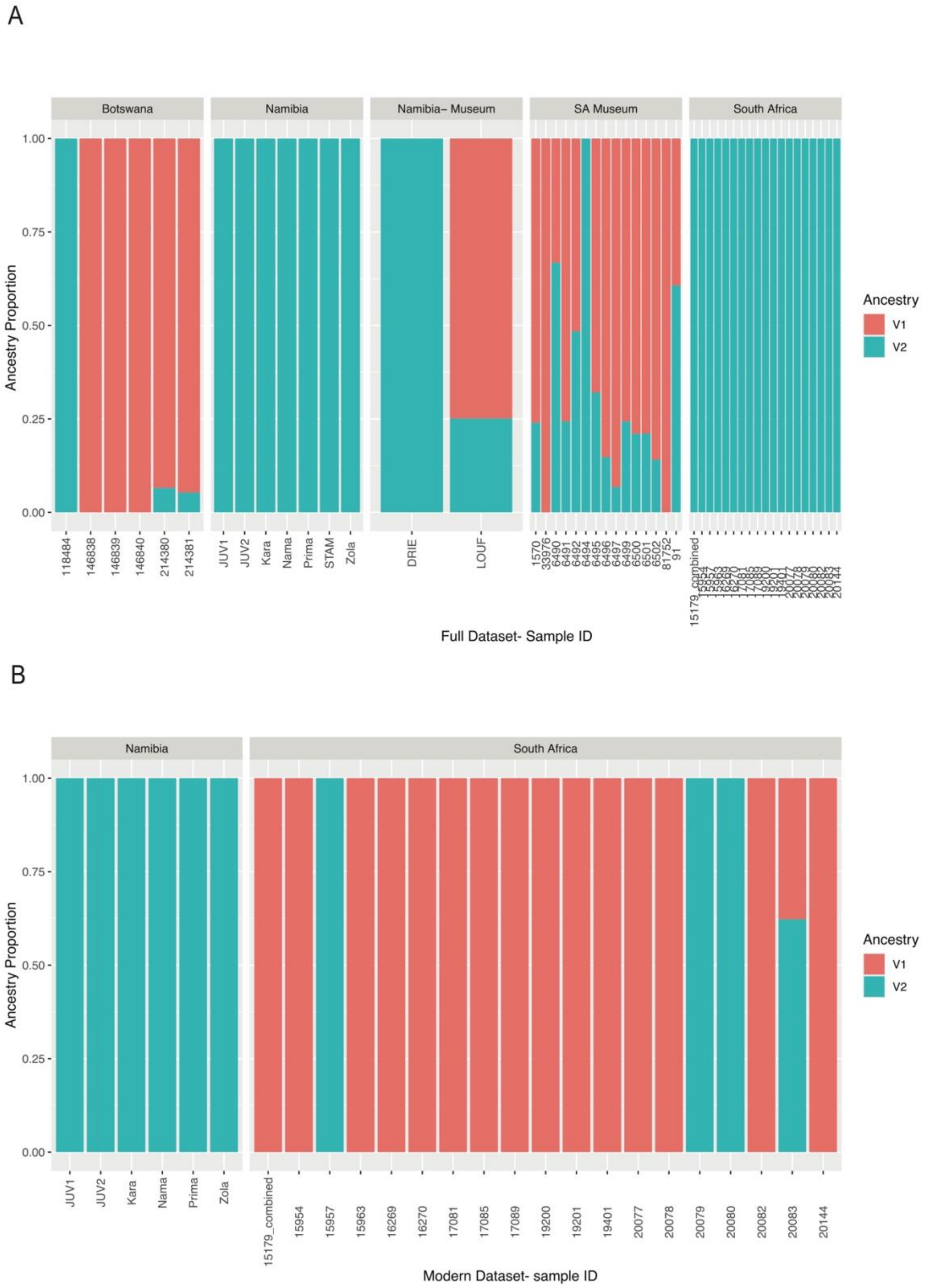
A
Botswana
Namibia
Namibia– Museum
SA Museum
South Africa
Ancestry Proportion
1.00
0.75
0.50
0.25
0.00
118484
146838
146839
146840
214380
214381
JUV1
JUV2
Kara
Nama
Prima
STAM
Zola
DRIE
LOUF
1570
33976
6490
6491
6492
6494
6495
6496
6497
6499
6500
6501
6502
81752
91
15179_combined
Ancestry
V1
V2
Full Dataset- Sample ID
B
Namibia
South Africa
Ancestry Proportion
1.00
0.75
0.50
0.25
0.00
JUV1
JUV2
Kara
Nama
Prima
Zola
15179_combined
15954
15957
15963
16269
16270
17081
17085
17089
19200
19201
19401
20077
20078
20079
20080
20082
20083
20144
Ancestry
V1
V2
Modern Dataset- sample ID

**Figure S3.** ADMIXTURE plot results for an analysis run with K=2. Each column represents the estimated fraction of the genome corresponding to each ancestry group in an individual. In **A)**, analysis reflects results from using the entire dataset. Samples are grouped by geographic origin with 'museum' samples noted in the header. In **B)**, results of the analysis using only modern samples are shown.

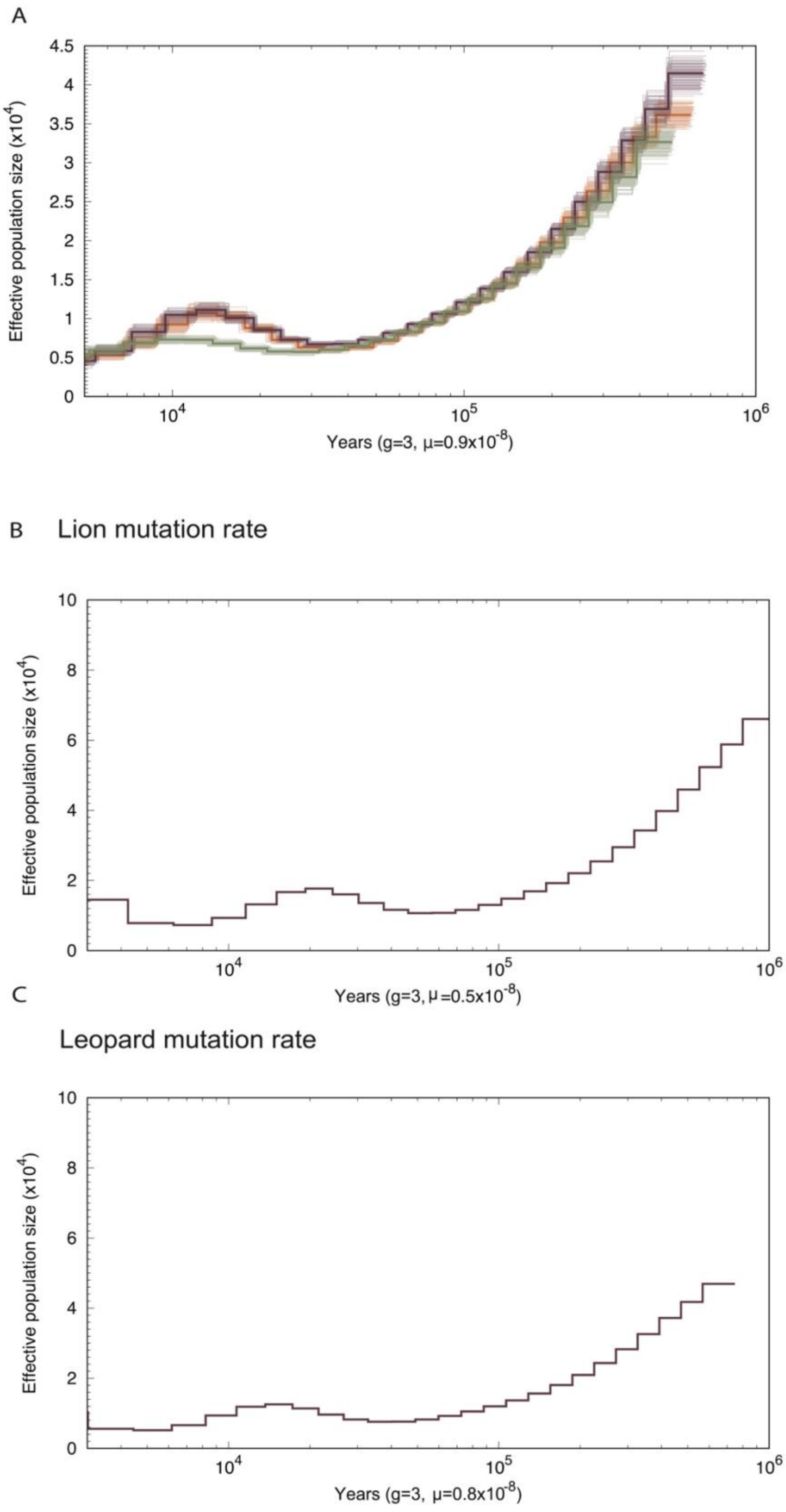
A
Effective population size (x10^4)
Years (g=3, μ=0.9x10^-8)
B Lion mutation rate
Effective population size (x10^4)
Years (g=3, μ=0.5x10^-8)
C
Leopard mutation rate
Effective population size (x10^4)
Years (g=3, μ=0.8x10^-8)

**Figure S4.** Effective population size ($N_e$) estimates reconstructed using the Pairwise Sequential Markovian Coalescent (Li & Durbin 2011) with 100 bootstrap replicates for each sample shown in lighter lines with colors corresponding to the country (Botswana – eggplant, Namibia – olive, and South Africa – orange). **A)** The South African sample appears shifted along the x-axis in comparison to the other two samples. We suspected that this may be due to coverage differences between samples; although we aimed for the same coverage across individuals, chance variation from pooling and filtering resulted in lower coverage for the South African sample (7X compared to >10X). To account for this difference, we downsampled the Botswana and Namibia samples to match the coverage of the South Africa sample. This resulted in concordant patterns across populations; these results are presented in main text Figure 2. **B & C)** We wanted to directly compare inferred historical population sizes in black-footed cats to other felid species. To do so, we reanalyzed our highest coverage individual using the mutation rates implemented in previous analyses of other Felids. Specifically, we reanalyzed data using **B)** the lion mutation rate of $5.4 \times 10^{-9}$ and **C)** the leopard mutation rate of $7.5 \times 10^{-9}$. We found that our results were qualitatively unchanged, with black-footed cats consistently inferred to have smaller effective population size compared to their respective counterparts.

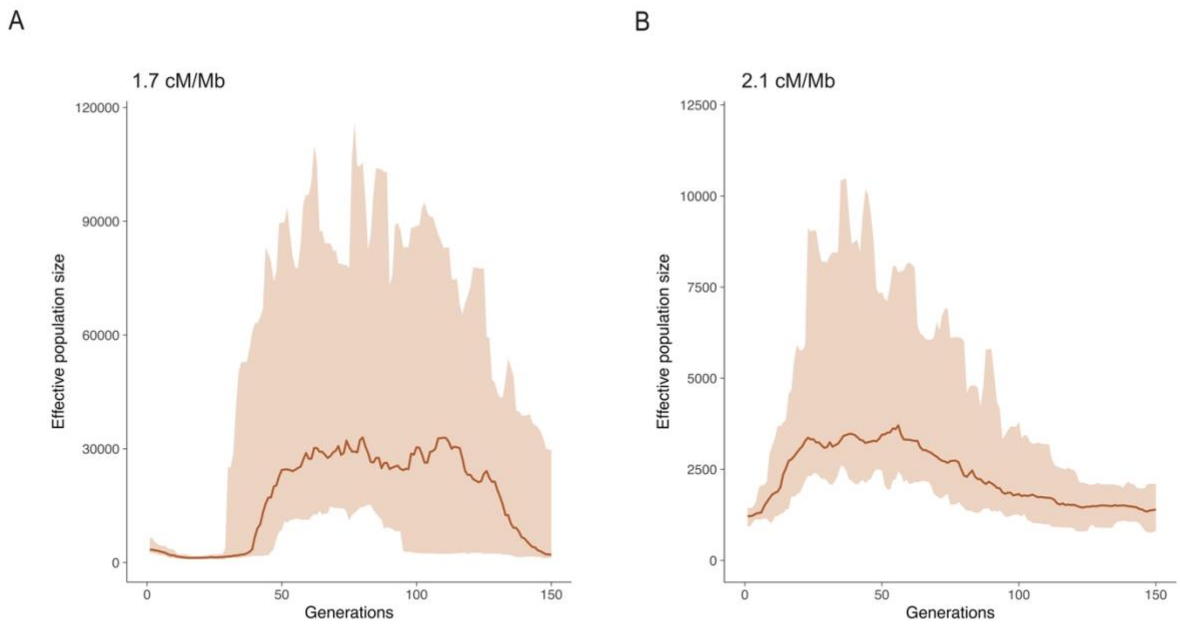


**Figure S5.** GONE analysis results intended to capture recent demographic history. The orange line shows the median estimates as a function of time. To produce confidence intervals, we ran 40 replicates. The shaded orange area shows the 95% confidence interval. GONE was run on default parameters with a modified recombination rate of **A)** 1.7 cM/Mb and **B)** 2.1 cM/Mb. In the main text figure (Figure 4B), we used the domestic cat recombination rate (1.9 cM/Mb), since recombination rate data is not available for black-footed cats. Note that estimates of effective population size varied dramatically as a function of the user-defined recombination rate.

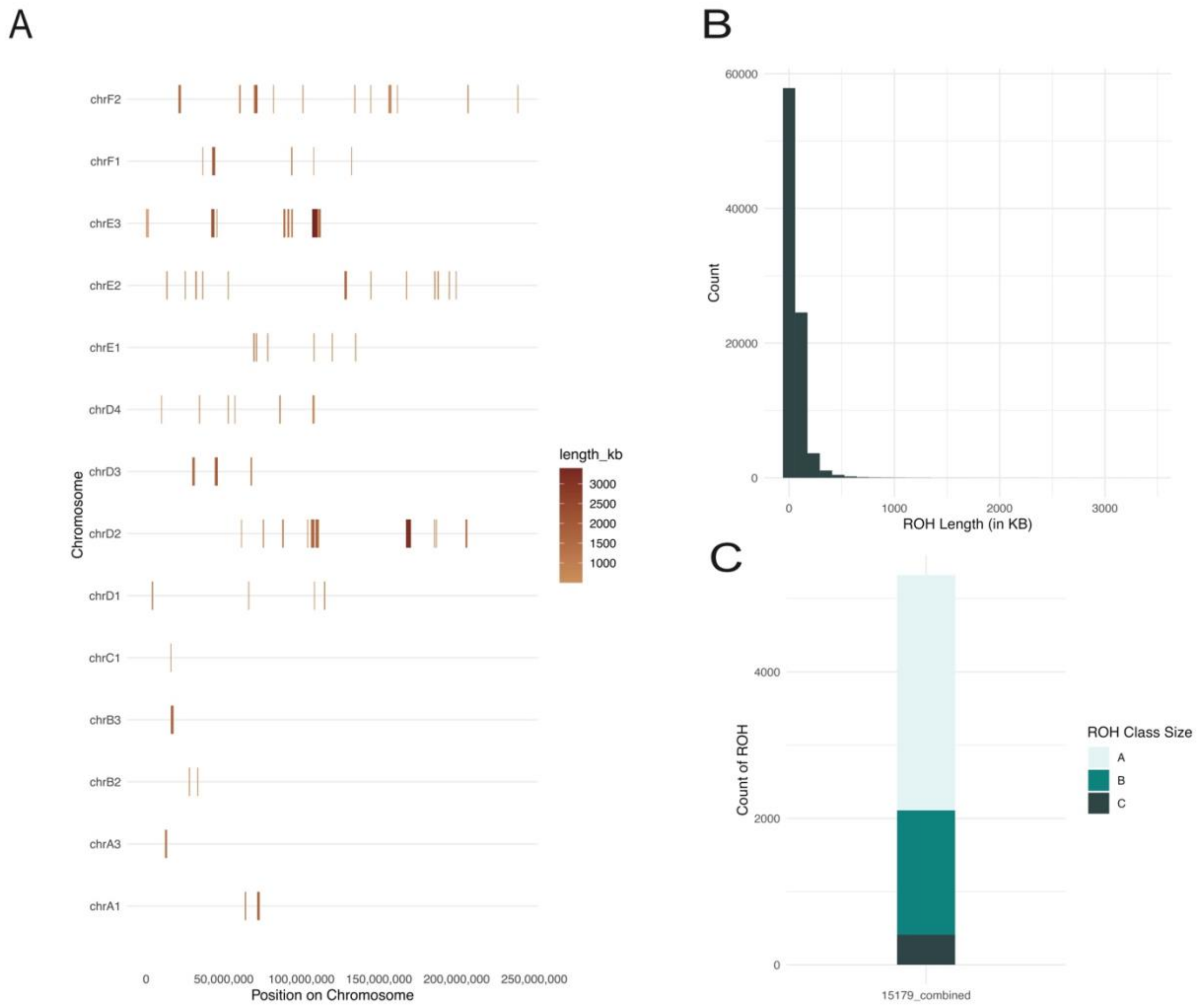


**Figure S6.** Runs of homozygosity (ROH) analysis inferred using GARLIC for one sample (15963) with 8X coverage. **A**) The Red-Blue color gradient shows the size of the inferred location and size of the ROH tract, with size increasing over the transition from light to dark. Only ROH tracts greater than 500 kb in length are plotted here. We also individually plotted counts of ROH tracts and size class to investigate ROH length. We found similar trends across samples; a representative individual is plotted here (015179). **B**) Histogram of detected ROH tracts as a function of ROH length. **C)** Histogram of ROH lengths broken into size class. Moving from shortest size class (A) to largest size class (C).

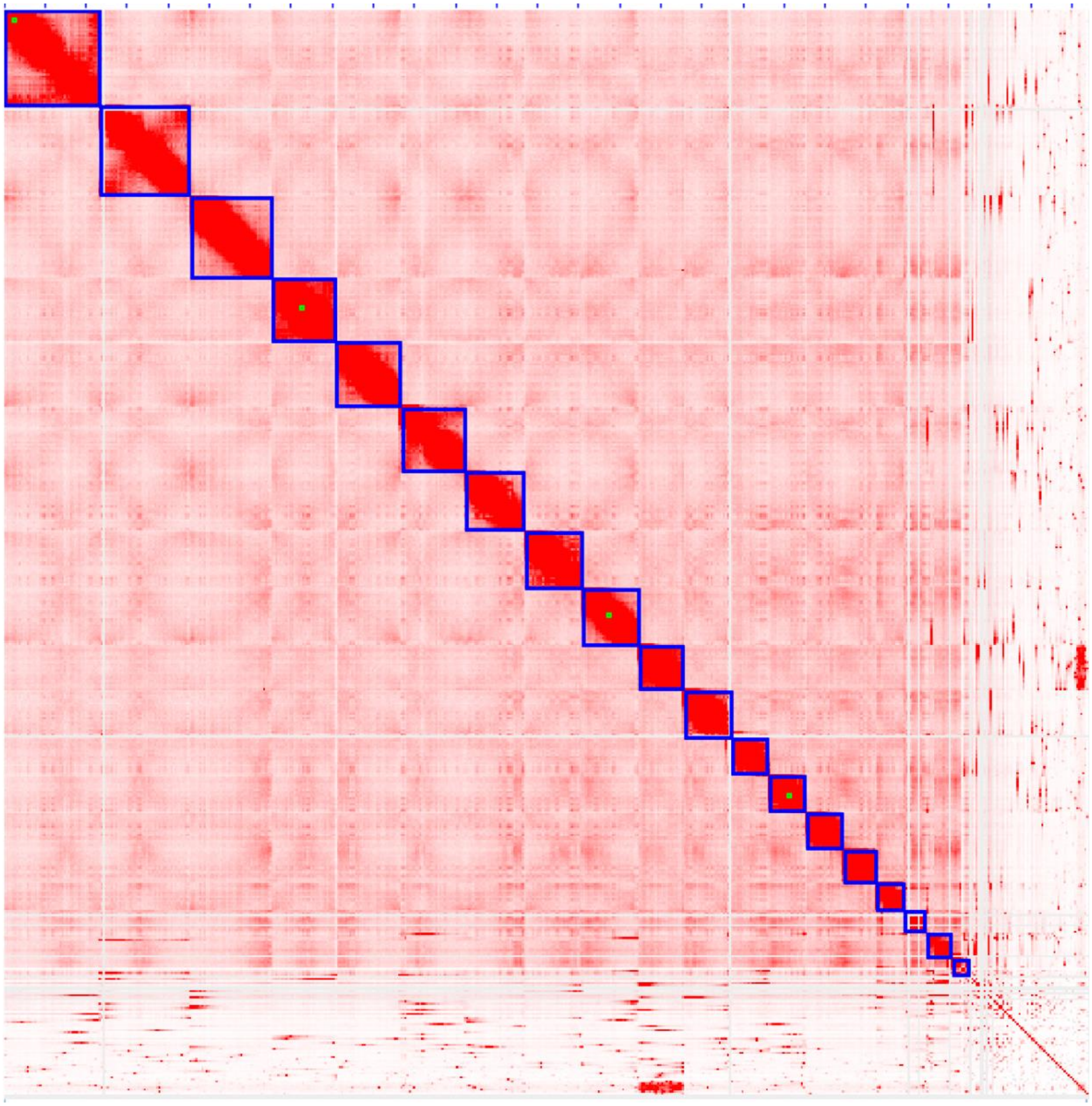

**Figure S7.** Hi-C interaction matrix of the black-footed cat, *Felis nigripes*. Genome-wide chromatin interaction map across 19 chromosomes. Red scale indicates contact interaction with deeper red color representing stronger evidence of contact. The horizontal and vertical axis represent the genomic position. Squares outlined in blue represent the highest intensity contacts and correspond to the 19 black-footed cat chromosomes.

**Supplementary Tables**

**Table S1**: Sample collection, mapping, and quality information. **Sample ID** lists sample name. **Library approach** refers to the library preparation protocol used for the samples (see main text methods). **Origin (Country)** identifies the country where the individual's biological sample was collected. **Origin (Locality)** identifies locality name of where the biological sample was collected. Individuals with missing locality data were noted with 'UNKNOWN' in the cell. The last three columns the contain number of sequence reads for each sample (**# of Reads**), the sample's sequence depth **(Sequence Depth)**, and the percentage of reads that mapped to our reference genome **(% of Mapped Reads**).

| Sample ID | Library Prep Approach | Origin (Country) | Origin (Locality) | # of Reads | Sequence Depth | % Mapped Reads |
|---|---|---|---|---|---|---|
| KB 015957 | Non-preserved | South Africa | Benfontein | 57,142,089 | 6.09221 | 96 |
| KB 16269 | Non-preserved | South Africa | Benfontein | 52,663,554 | 5.6119 | 96 |
| KB 19201 | Non-preserved | South Africa | UNKNOWN | 47,095,934 | 4.99053 | 94 |
| KB 15956 | Non-preserved | South Africa | Benfontein | 126,395,156 | 13.2327 | 99 |
| KB 15963 | Non-preserved | South Africa | Benfontein | 303,768,014 | 32.1175 | 99 |
| KB 15179 | Non-preserved | South Africa | Benfontein | 232,190,018 | 24.7067 | 99 |
| KB 19200 | Non-preserved | South Africa | UNKNOWN | 229,397,772 | 23.4921 | 99 |
| KB 20144 | Non-preserved | South Africa | UNKNOWN | 153,617,071 | 16.3285 | 99 |
| KB 15954 | Non-preserved | South Africa | Benfontein | 85,267,687 | 9.08205 | 98 |
| KB 16270 | Non-preserved | South Africa | Benfontein | 83,477,506 | 8.87537 | 98 |

| | | | | | | |
|---|---|---|---|---|---|---|
| KB 17081 | Non-preserved | South Africa | Benfontein | 111,402,021 | 11.7752 | 99 |
| KB 17085 | Non-preserved | South Africa | De Aar | 90,133,118 | 9.30752 | 98 |
| KB 17089 | Non-preserved | South Africa | Benfontein | 125,122,473 | 12.9052 | 99 |
| KB 19401 | Non-preserved | South Africa | UNKNOWN | 90,678,482 | 9.55013 | 98 |
| KB 20077 | Non-preserved | South Africa | UNKNOWN | 91,604,104 | 9.74222 | 98 |
| KB 20078 | Non-preserved | South Africa | UNKNOWN | 120,780,194 | 12.844 | 99 |
| KB 20079 | Non-preserved | South Africa | UNKNOWN | 82,622,533 | 8.60365 | 97 |
| KB 20080 | Non-preserved | South Africa | UNKNOWN | 74,315,571 | 7.87882 | 97 |
| KB 20082 | Non-preserved | South Africa | UNKNOWN | 82,867,334 | 8.69891 | 98 |
| KB 20083 | Non-preserved | South Africa | UNKNOWN | 109,303,346 | 11.2534 | 98 |
| OR 5264 | Non-preserved | South Africa | UNKNOWN | 108,592,588 | 11.58 | 99 |
| 6502 | Preserved | South Africa | Uppington | 76,648,264 | 1.04094 | 52 |
| 6501 | Preserved | South Africa | Uppington | 55,066,103 | 1.0441 | 58 |
| 6497 | Preserved | South Africa | Hopetown | 82,915,448 | 1.37559 | 64 |
| 6495 | Preserved | South Africa | Kimberly | 46594265 | 1.17669 | 61 |
| 6494 | Preserved | South Africa | Coppertown | 463,340,726 | 9.99388 | 86 |
| 6492 | Preserved | South Africa | Setlagole | 375,324,219 | 7.71039 | 84 |
| 6490 | Preserved | South Africa | NA | 47,208,425 | 1.18355 | 60 |
| 91 | Preserved | South Africa | Gordonia | 387,393,204 | 7.1549 | 84 |
| BFC-214381 | Preserved | South Africa | Thornfloof | 89,589,107 | 1.11022 | 57 |

| | | | | | | |
|---|---|---|---|---|---|---|
| BFC-33976 | Preserved | South Africa | Transvaal | 102,603,089 | 1.57515 | 68 |
| BFC-81752 | Preserved | South Africa | Kroonstad | 89,538,552 | 1.11483 | 54 |
| BFC-JUV1 | Non-preserved | Namibia | Grünau | 126,049,656 | 2.51322 | 78 |
| BFC-JUV2 | Non-preserved | Namibia | Grünau | 145,653,105 | 3.20774 | 81 |
| BFC-Kara | Non-preserved | Namibia | Grünau | 128,126,409 | 2.80737 | 79 |
| BFC-Nama | Non-preserved | Namibia | Grünau | 146,908,603 | 3.35932 | 81 |
| BFC-Prima | Non-preserved | Namibia | Grünau | 112,304,230 | 2.53537 | 78 |
| BFC-Zola | Non-preserved | Namibia | Grünau | 109,082,911 | 2.06656 | 75 |
| BFC-STAM | Preserved | Namibia | Stampriet-aranos road | 112,435,626 | 3.22172 | 80 |
| BFC-DRIE | Preserved | Namibia | Grünao | 285,136,135 | 7.90913 | 85 |
| BFC-118484 | Preserved | Botswana | Tstestin | 363,653,340 | 10.2557 | 86 |
| BFC-146838 | Preserved | Botswana | Molepolole | 85,454,683 | 1.19463 | 60 |
| BFC-146839 | Preserved | Botswana | Molepolole | 93,847,507 | 1.58993 | 67 |
| BFC-146840 | Preserved | Botswana | Molepolole | 106,311,813 | 1.71304 | 69 |

**Table S2**: Assembly statistics for various builds of the black-footed cat genome. The **Primary Hifiasm** column refers to the first stage of assembly where the draft genome was assembled with PacBio data and joined with Hi-C data using Hifiasm with no scaffolding or filtering. The second column, **Primary YaHS Hi-C**, is the second assembly stage post scaffolding and filtering using YaHS v1.1 (Zhou et al. 2023). The third column, **Post join & clean (BFC_Final)**, is the final genome assembly after joining and filtering (see main text methods) used in this study. The fourth column, **BFC-DNAZoo**, refers to the draft genome available on the DNAZoo website. The final column, **Felcat9**, consists of assembly statistics from the Felcat9 domestic cat genome assembly available on NCBI.

| | **Primary Hifiasm** | **Primary YaHS Hi-C** | **Post join & clean (BFC_Final)** | **BFC-DNAZoo** | **Felcat9** |
|---|---|---|---|---|---|
| **Length** | 2,642,259,949 | 2,642,793,549 | 2,642,807,549 | 2,454,753,549 | 2,521,710,482 |
| **# of scaffolds** | NA | 2,526 | 2,456 | 226,988 | 4,507 |
| **Scaffold N50** | NA | 139,171,707 | 142,612,561 | 139,656,774 | 14,9751,809 |
| **Scaffold L90** | NA | 64 | 19 | 18 | 16 |
| **#N** | NA | 533,600 | 547,600 | 16,838,500 | 45,197,951 |
| **# of contigs** | 5,115 | 5,194 | 5,194 | 286,461 | 4,800 |
| **Contig N50** | 1,475,238 | 1,469,769 | 1,469,769 | 50,735 | 41,915,695 |
| **Contig L90** | 2,090 | 2,117 | 2,117 | 50,172 | 71 |

**Table S3:** BUSCO statistics for various *Felis* genome assemblies based on the mammalia_odb10 dataset. The row names refer to the different genome assemblies being compared including the genome assembly in this study (**BFC_Final**), the publicly available DNAZoo assembly (**BFC-DNAZoo**), and an outgroup, the jungle cat, available on NCBI (***Felis chaus***). The column names refer to the subset of conserved orthologous genes analyzed for each genome assembly. Listed in each column is the percentage of the conserved orthologous genes within the focal genome assembly based on analysis with the complete set (**Complete**), the single-copy set (**Single-copy**), duplicated genes (**Duplicated**), fragmented genes (**Fragmented**), and the proportion of genes missing from the set (**Missing**).

| | **Complete** | **Single-copy** | **Duplicated** | **Fragmented** | **Missing** |
|---|---|---|---|---|---|
| **BFC_Final** | 94.3% | 92.3% | 2.0% | 1.4% | 4.3% |
| **BFC-DNAZoo** | 89.6% | 89.1% | 0.5% | 3.9% | 6.5% |
| ***Felis chaus*** | 95.8% | 95.2% | 0.6% | 1.0% | 3.2% |

**Table S4:** BUSCO statistics for various reference assemblies of *Felis* using the carnivora_odb10 dataset. Row names indicate the names of the different genome assemblies whose BUSCO statistics are being compared including this study's genome assembly (**BFC_Final**), the DNAZoo black-footed cat assembly (**BFC-DNAZoo**), the NCBI domestic cat genome (**Felcat9**), and an outgroup, the jungle cat, available on NCBI (***Felis chaus***). The column names refer to the subset of conserved orthologous genes analyzed for each genome assembly. Listed in each column is the percentage of the conserved orthologous genes within the focal genome assembly based on analysis with the complete set (**Complete**), the single-copy set (**Single-copy**), duplicated genes (**Duplicated**), fragmented genes (**Fragmented**), and the proportion of genes missing from the set (**Missing**).

| | Complete | Single-copy | Duplicated | Fragmented | Missing |
|---|---|---|---|---|---|
| BFC_Final | 93.5% | 91.4% | 2.1% | 1.2% | 5.3% |
| BFC-DNAZoo | 91.2% | 90.5% | 0.7% | 2.1% | 6.7% |
| Felcat9 | 95.1% | 94.2% | 0.9% | 0.9% | 4.0% |
| ***Felis chaus*** | 95.2% | 94.4% | 0.8% | 0.9% | 3.9% |

**Table S5.** For several analysis, we generated downsampled BAM files with the goal of understanding the role of technical factors in our results. Information on the results of downsampling are listed here. **Sample ID** indicates sample name. Samples with >2X coverage in our dataset were downsampled to ~1X coverage to generate similar coverage for downstream analyses. The resulting coverage of these samples is noted in **Down-sample BAM Coverage**. The respective columns name the analyses that used these BAM files in our study. X indicates analyses where down-sampled BAM files were used. BAM files with duplicated down-sample files were noted. The + symbol marks BAM files that were downsampled to two different target coverage levels: once to generate a ~1X file for population genetic comparisons and once to generate 7X coverage for analyses where ultra-low coverage data cannot be used. The * symbol indicates BAM files that were downsampled to 7X for a distinct set of analyses

| Sample ID | Down-sample BAM Coverage | PSMC | Pixy | Dxy | PCA | NGSadmix |
|---|---|---|---|---|---|---|
| **KB 015957** | 1.2 | | | | X | X |
| **KB 16269** | 1.1 | | | | X | X |
| **KB 19201** | 1 | | | | X | X |
| **KB 15956** | 1.3 | | | | X | X |
| **KB 15963** | 1.6+ | | | | X | X |
| **KB 15179** | 1.2 | | | | X | X |
| **KB 19200** | 1.2 | | | | X | X |
| **KB 20144** | 1.6 | | | | X | X |
| **KB 15954** | 1.4 | | | | X | X |
| **KB 16270** | 1.3 | | | | X | X |
| **KB 17081** | 1.2 | | | | X | X |
| **KB 17085** | 1.4 | | | | X | X |
| **KB 17089** | 1.3 | | | | X | X |
| **KB 19401** | 1 | | | | X | X |
| **KB 20077** | 1.2 | | | | X | X |
| **KB 20078** | 1.3 | | | | X | X |
| **KB 20079** | 1.3 | | | | X | X |
| **KB 20080** | 1.2 | | | | X | X |
| **KB 20082** | 1.3 | | | | X | X |
| **KB 20083** | 1.1 | | | | X | X |
| **OR 5264** | 1.2 | | | | X | X |
| **6502** | 1 | | | | X | X |
| **6501** | 1 | | | | X | X |
| **6497** | 1 | | | | X | X |
| **6495** | 1 | | | | X | X |
| **6494** | 1+, 7.5 | X* | X* | | X | X |
| **6492** | 1.2 | | | | X | X |
| **6490** | 1 | | | | X | X |
| **91** | 1.4 | | | | X | X |
| **BFC-214381** | 1 | | | | X | X |
| **BFC-33976** | 2 | | | | X | X |

| | | | | | | |
|---|---|---|---|---|---|---|
| **BFC-81752** | 1 | | | | X | X |
| **BFC-JUV1** | 1.3 | | | | X | X |
| **BFC-JUV2** | 1.3 | | | | X | X |
| **BFC-Kara** | 1.4 | | | | X | X |
| **BFC-Nama** | 1.3 | | | | X | X |
| **BFC-Prima** | 1.3 | | | | X | X |
| **BFC-Zola** | 1.03784 | | | | | |
| **BFC-STAM** | 1.6 | | | | X | X |
| **BFC-DRIE** | 1.2+, 7.9 | X* | X* | | X | X |
| **BFC-118484** | 1.2+, 7.7 | X* | X* | | X | X |
| **BFC-146838** | 1.7 | | | | X | X |
| **BFC-146839** | 2 | | | | X | X |
| **BFC-146840** | 2 | | | | X | X |
